# A novel DNA repair protein, N-Myc downstream regulated gene 1 (NDRG1), links stromal tumour microenvironment to chemoresistance

**DOI:** 10.1101/2025.01.22.634323

**Authors:** Nina Kozlova, Kayla A Cruz, Hanna M Doh, Antoine A Ruzette, Nicholas A Willis, Su Min Hong, Raul S Gonzalez, Monika Vyas, Laura M Selfors, Stephan Dreyer, Rosie Upstill-Goddard, Kerrie L Faia, Steve Wenglowsky, Josh Close, Alica K. Beutel, Zeljka Jutric, Michael U J Oliphant, Byanjana Thapa, Martin S Taylor, Venla Mustonen, Pradeep Mangalath, Christopher J Halbrook, Joseph E Grossman, Rosa F Hwang, John G Clohessy, Salla Ruskamo, Petri Kursula, Boryana Petrova, Naama Kanarek, Philip A Cole, David K Chang, Simon F Nørrelykke, Ralph Scully, Taru Muranen

## Abstract

In pancreatic ductal adenocarcinoma cancer (PDAC) drug resistance is a severe clinical problem and patients relapse within a few months after receiving the standard-of-care chemotherapy. One contributing factor to treatment resistance is the desmoplastic nature of PDAC; the tumours are surrounded by thick layers of stroma composing up to 90% of the tumour mass. This stroma, which is mostly comprised of extracellular matrix (ECM) proteins, is secreted by cancer-associated fibroblasts (CAFs) residing in the tumour microenvironment. However, the mechanistic basis by which the tumour stroma directly contributes to chemoresistance remains unclear. Here, we show that CAF-secreted ECM proteins induce chemoresistance by blunting chemotherapy-induced DNA damage. Mechanistically, we identify N-myc downstream regulated gene 1 (NDRG1) as a key protein required for stroma-induced chemoresistance that responds to signals from the ECM and adhesion receptors. We further show that NDRG1 is a novel DNA repair protein that physically interacts with replication forks, maintains DNA replication and functions to resolve stalled forks caused by chemotherapy. More specifically, NDRG1 reduces R-loops, RNA-DNA hybrids that are known to cause genomic instability. R-loops occur during replication-transcription conflicts in S-phase and after chemotherapy treatments, thus posing a major threat to normal replication fork homeostasis. We identify NDRG1 as highly expressed in PDAC tumours, and its high expression correlates with chemoresistance and poor disease-specific survival. Importantly, knock-out of NDRG1 or inhibition of its phosphorylation restores chemotherapy-induced DNA damage and resensitizes tumour cells to treatment. In conclusion, our data reveal an unexpected role for CAF-secreted ECM proteins in enhancing DNA repair via NDRG1, a novel DNA repair protein, directly linking tumour stroma to replication fork homeostasis and R-loop biology, with important therapeutic implications for restoring DNA damage response pathways in pancreatic cancer.

**Summary paragraph:** Drug resistance is a severe clinical problem in stroma-rich tumours, such as pancreatic ductal adenocarcinoma (PDAC), and patients often relapse within a few months on chemotherapy^1–9^. The stroma, comprised of extracellular matrix (ECM) proteins, is secreted by cancer-associated fibroblasts (CAFs) residing in the tumour microenvironment^10–13^. Prior work show that ECM proteins provide survival benefits to cancer cells^14,15^. However, the precise role of CAF-secreted ECM in resistance to DNA damaging chemotherapies remains poorly understood. Here, we link ECM proteins to chemoresistance by enhanced DNA damage repair (DDR). Mechanistically, we identify N-myc downstream-regulated gene 1 (NDRG1) as a key effector downstream of ECM and the integrin-Src-SGK1-signalling axis that mediates enhanced DDR. We show that *NDRG1* loss, mutation of conserved His194, or inhibition of NDRG1 phosphorylation by SGK1 lead to replication fork stalling, increased R-loops, and higher transcription-replication conflicts, resulting in genomic instability and sensitivity to chemotherapies. Our analysis of PDAC patient cohorts^16^ found that high NDRG1 expression correlates with chemoresistance and poor patient survival. In conclusion, we uncover an unexpected role for CAF-secreted ECM proteins in promoting therapeutic resistance by enhancing DDR and establish NDRG1 as a novel DNA repair protein directly linking tumour stroma to DDR.

## Introduction

Pancreatic ductal adenocarcinoma (PDAC) is one of the most lethal cancers with a 13% 5-year survival rate. It is also one of the few cancers where the incidence in the U.S. is still increasing. Projections show that about 66,440 new cases will be diagnosed in the U.S. in 2024 and 51,750 patients will die of their disease (American Cancer Society, Cancer statistics 2024). The standard-of-care in PDAC is chemotherapy, with combination treatment FOLFIRINOX (Folinic acid-Fluorouracil-Irinotecan-Oxaliplatin) being the most effective, but it also has toxic side effects. In the event of intolerable toxicity, patients switch to gemcitabine (GEM) as a single agent, or in combination with Albumin-bound paclitaxel (Abraxane, GA) depending on patient-specific factors^1^. Most of these chemotherapies work by inducing DNA damage or stalling DNA replication. Importantly, PDAC tumours usually develop chemoresistance rapidly, resulting in patient death. Therefore, a critical need exists to understand what drives the emergence of chemoresistance in PDAC and to develop more effective treatment strategies to improve patient outcomes.

Resistance to chemotherapies can occur through a variety of mechanisms that include, but are not limited to, alterations in the drug availability, drug metabolism, enhanced DNA repair, and upregulation of pro-survival signalling in cancer cells^17^. In PDAC, one of the major factors contributing to chemoresistance is the dense stroma. Primarily, the stroma poses a physical barrier that reduces the diffusion of therapeutic agents and promotes the degradation of drugs by stromal enzymes^7^. In addition, high extracellular matrix (ECM) content provides various survival benefits for cancer cells. It has been shown that human PDAC tumours are low in glucose, upper glycolytic intermediates, glutamine and serine, rendering the pancreatic tumour environment nutrient-starved and dependent on enhanced micropinocytosis for nutrient scavenging^18^. Our previous work showed that in a starved environment cells uptake ECM proteins as a source of amino acids^14^. Similarly, under restrictive nutrient conditions, PDAC cells were found to uptake stroma-secreted alanine^19^ and collagen-derived proline^15^ for survival.

Cancer-associated fibroblasts (CAFs) represent a highly abundant and heterogenous cell population^12,13^ that are responsible for the abundant secretion of ECM proteins observed in PDAC^9^. These CAFs arise from various sources including resident fibroblasts, mesenchymal stem cells, or other cell types like pancreatic stellate cells (PSCs). Activation of PSCs and CAFs can occur in response to stresses like inflammation, alcohol metabolites, growth factors (e.g. PDGF, TGF-α, TGF-β) or hypoxia^20^. Once activated, PSCs transdifferentiate into alpha-smooth muscle actin (α-SMA) expressing CAFs that secrete large amounts of ECM into the tumour microenvironment (TME).

Since we previously found a significant increase in ECM secretion by starved primary human fibroblasts^14^, we hypothesized that the nutrient-poor environment in the pancreatic tumours might activate the pancreatic CAFs and lead to enhanced ECM secretion. While stroma-rich tumours are known to be highly resistant to anti-cancer treatments^5^, a mechanistic basis by which the tumour stroma directly contributes to DNA repair that in turn can lead to chemoresistance is not known. Therefore, this work aimed to identify a mechanistic connection between the high abundance of ECM in the PDAC TME and how this might stimulate chemoresistance in the tumour cells.

## Results

### Nutrient deprivation activates pancreatic fibroblasts, leading to increased secretion of ECM proteins and increased chemoresistance

Given the nutrient-poor conditions and desmoplasia frequently observed in PDAC, we wanted to assess how nutrient starvation influences patient-derived pancreatic CAFs (Extended Data Fig. 1a-b). When CAFs were serum and lipid-starved we observed a significant increase in ECM secretion (Fig. 1a, Extended Data Fig. 1c-d), similar to what has been observed under hypoxic conditions^21^. Pancreatic CAFs have been previously linked to tumour growth and treatment resistance^12^. Therefore, we assessed whether the CAF-conditioned media (CAF-CM) harvested from starved fibroblasts would also induce treatment resistance. Indeed, CAF-CM provided a proliferative advantage to cancer cells (Extended Data Fig. 1e-f) and stimulated resistance against multiple chemotherapies used in the treatment of PDAC (Extended Data Fig. 1g-j). Most chemotherapies work by inflicting DNA damage (e.g., gemcitabine, irinotecan (SN-38), 5-fluorouracil, oxaliplatin). Therefore, we wanted to determine whether CAF-CM from different CAF lines could stimulate chemoresistance in PDAC cells by reducing DNA damage. We analysed DNA damage by yH2AX staining of cells treated with CAF-CM vs. non-conditioned media (NC) in the presence of chemotherapies. CAF-CM significantly reduced chemotherapy-nduced DNA damage (Fig. 1b-c, Extended Data Fig. 1k-o). We further validated these findings by using an alkaline Comet assay that measures levels of single stranded DNA breaks on a single cell level, and used Olive Tail Moment (OTM) as a parameter to measure the amount of damaged DNA^22^. This assay further confirmed that CAF-CM significantly reduces chemotherapy-induced DNA damage (Fig. 1d).

**Fig. 1.**
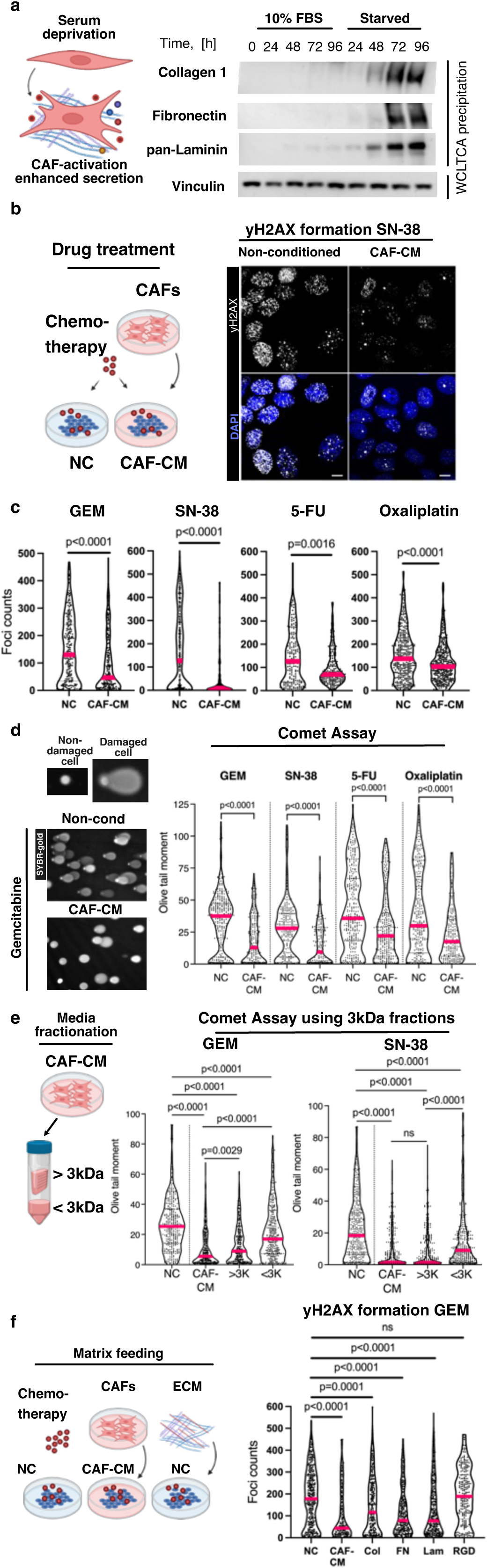
Conditioned media from starved CAFs increases chemoresistance in pancreatic cancer cells. **a**, Pancreatic CAFs were grown in DMEM containing 10% FBS or 0% FBS and media and cells were harvested at indicated times. CAF medium was TCA precipitated and immunoblotted for Collagen 1, Fibronectin, and pan-Laminin. Whole cell lysates (WCL) of the CAFs were probed for Vinculin. TCA loading was normalized to the WCL protein amount. **b-c,** yH2AX foci were used to assess DNA damage. SW1990 were treated with 1 µM gemcitabine (GEM), 0.5 µM SN-38, 100 µM 5-FU, or 50 µM oxaliplatin for 24h in non-conditioned media (NC) or CAF-conditioned media (CAF-CM). Cells were fixed and immunostained for yH2AX (white) and DAPI (blue) followed by image acquisition. Images were quantified by CellProfiler; data in (**c**) are shown as foci counts per single nucleus. >100 nuclei were analysed per condition. Mann-Whitney test was applied, magenta bars represent median. Representative images in (**b**) display yH2AX foci formation upon SN-38 treatment. Scale bar: 10 µm. **d**, SW1990 were analysed for DNA damage by the alkaline comet assay. Cells were treated with indicated chemotherapies as in (**b-c**) for 24h in NC or CAF-CM, plated on Comet assay slides, DNA was run under alkaline conditions and stained with SYBR-gold. Non-damaged DNA appears as a dense dot, while the loops of the damaged DNA lose their supercoiling and form comet tails, a representative image is shown for GEM. Images were quantified using Comet Score software. Olive Tail Moment (OTM), reflecting the intensity of the comet tail relative to the head, was used as a metric for the level of DNA damage. Higher OTM signifies higher levels of DNA damage. Data were analysed in Graphpad Prism8 and Mann-Whitney test was used to establish significance. Magenta lines represent median, at least 200 cells were analysed for each treatment condition. **e**, Proteins were filtered from CAF-CM using 3kDa cut-off spin filters. SW1990 cells were treated with 1 µM GEM or 0.5 µM SN-38 in the presence of NC, CAF-CM, or with filtered CAF-CM (<3KDA proteins or <3kDa proteins as indicated). Comet assay was used to analyse DNA damage, OTM values were plotted and Mann-Whitney test was used to examine significance (*n*≥200). Magenta lines represent median values. **f**, SW1990 cells were plated on coverslips, serum starved overnight, and indicated ECM proteins (Col: Collagen 1, FN: Fibronectin, Lam: Laminin-5), RGD peptides, NC, or CAF-CM were added to the cells along with 0.5 µM GEM for 24h. Cells were fixed and stained for yH2AX and DAPI, and the images were quantified for yH2AX foci/nucleus by CellProfiler software. Mann-Whitney test was applied, magenta bars represent median. N=3.

Several studies have described that CAF-CM provides growth and survival advantages to PDAC tumour cells through multiple mechanisms, including metabolic reprograming, and through secretion of deoxycytidine that competes with gemcitabine for the incorporation into the nascent DNA^23–27^. However, our data suggest that in addition to gemcitabine, CAFs protect PDAC cells against irinotecan, oxaliplatin, and 5-FU, pointing to a broader mechanism of CAF-mediated chemoresistance. To assess whether the protective effect of CAF-CM is reliant on deoxycytidine or other small metabolites, we filtered the CAF-CM through a 3 kDa filter. The eluted fraction (<3kDa) that contained metabolites provided a modest protection against gemcitabine as previously reported^23–27^. Interestingly, this fraction also provided modest protection against SN-38, which was unlikely to be mediated by secreted deoxycytidine, raising the possibility that other secreted metabolites may also play a role in chemoresistance (Fig. 1e, Extended Data Fig. 1p-s, and Extended Data Table 1). However, the retained fraction (>3kDa) provided a significantly stronger protection against gemcitabine and SN-38 (Fig. 1e, Extended Data Fig. 1p-s). The protective effect disappeared after boiling the CAF-CM (Extended Data Fig. 1p), further suggesting a protein-mediated resistance mechanism. The OTM values and the yH2AX foci of cells treated with >3kDa fraction were in most cases comparable to cells treated with non-fractionated CAF-CM, further suggesting a protein-mediated resistance mechanism. Overall, these data show that CAF-CM protects PDAC cells from multiple chemotherapies by decreasing DNA damage, most likely via a protein-related mechanism.

Because the CAFs are known to secrete high amounts of ECM proteins, we next wanted to assess whether secreted ECM proteins could be responsible for the increased tolerance towards DNA damaging agents. Purified ECM proteins (collagen 1, fibronectin, laminin) were added to tumour cells after which the cells were treated with DNA damaging agents for 24h (SN-38 or gemcitabine) and analysed for yH2AX foci. We observed fewer yH2AX foci in cells treated with purified ECM proteins, suggesting that multiple purified ECM proteins can reduce chemo-induced DNA damage (Fig. 1f, Extended Data Fig. 1t-v). Although all the tested ECM proteins provided protection, there was some variation in DNA damage response between the ECM proteins used, suggesting that multiple integrin heterodimers might provide protection towards DNA damaging agents. Indeed, analysis of the integrin expression of PDAC cell lines showed high heterogeneity, which was further modulated by the addition of CAF-CM (Extended Data Fig. 1w-x). RGD peptides alone were not effective, suggesting that larger or polymerized ECM proteins are required for the observed protection (Fig. 1f, Extended Data Fig. 1t-v).

### Phospho-N-myc downstream regulated gene 1 (NDRG1) is upregulated in PDAC cells in response to CAF-CM

To gain more mechanistic insight into how the tumour cells respond to CAF-CM and secreted ECM proteins, and to identify potential drivers of chemoresistance we performed a reverse-phase protein array (RPPA) that measures in a quantitative manner ∼304 proteins and phospho-proteins representing major signalling hubs and pathways in cancer^28–30^. We performed RPPA on three PDAC cell lines (BxPC-3, HPAC, SW1990) in the presence of gemcitabine, the most frequently used mono-therapy agent in PDAC. Multiple pro-survival proteins were up-regulated in the presence of CAF-CM at 72h (Fig. 2a-b). These included p-STAT3, a well-known protein that responds to CAF-CM, EGFR, and IGF1R. Not surprisingly ECM-responsive proteins such as p-Src and p-FAK, which respond to integrin activation, were also up-regulated in the tumour cells incubated with the CAF-CM. However, the top-scoring candidate in all the PDAC lines was phospho-NDRG1 T346 (Fig. 2a), with p-NDRG1 levels up-regulated at 72h, a timepoint at which we also observed higher total-NDRG1 levels (Fig. 2b). NDRG1 is a stress-response protein with a somewhat unclear role in cancer. While it is described to be a cell-migration suppressor in prostate and gastric cancer cell lines^31–33^, it is an indicator of poor prognosis for patients with glioma, inflammatory breast, gallbladder and liver cancer^34–38^. Since p-NDRG1 was highly up-regulated with CAF-CM, we sought to unravel the consequences of this observation in PDAC.

**Fig. 2.**
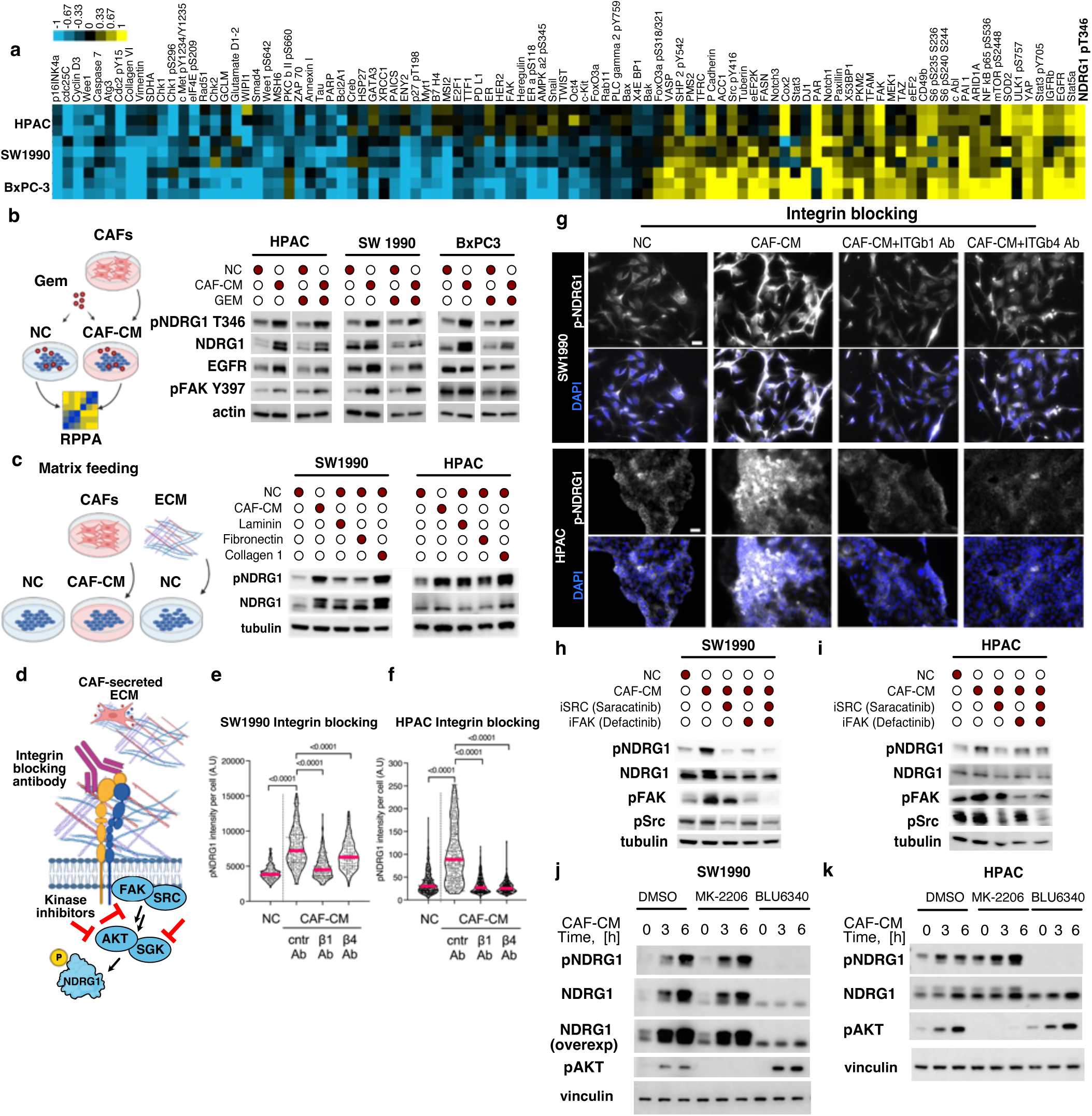
Phospho-NDRG1 is up-regulated by CAF-conditioned media and is phosphorylated via ECM-integrin-Src-FAK-SGK1-axis. **a,** Reverse Phase Protein Array (RPPA) was performed on three pancreatic cancer cell lines (HPAC, SW1990, BxPC-3), probing for 304 phospho- and total proteins. The cells were treated with 1 µM gemcitabine (GEM) for 72h in NC or CAF-CM. The data are shown as CAF-CM media normalized over NC media. Only significantly (p<0.05) changed proteins are shown. Yellow indicates proteins that were most significantly up-regulated in the CAF-CM and blue indicates down-regulated proteins (heat-map: log2 fold change). **b**, Western blot validation of the RPPA. Cells were treated with NC media or CAF-CM +/-1 µM GEM for 72h, media were changed daily, and cells were harvested 1h after the last media change. Gels were run at least thrice with similar results and probed for p-NDRG1 (T346), total NDRG1, EGFR, p-FAK (Y397), and actin. **c**, Purified ECM proteins (Laminin-5, Fibronectin, or Collagen 1) were added to serum starved HPAC or SW1990 cells for 3h, NC and CAF-CM media were used as controls. Cells were lysed and analysed for p-NDRG1, total NDRG1, and tubulin. **d**, General scheme of ECM-triggered signalling events, and pharmacological inhibition approaches used in experiments **e-k**. **e-g**, SW1990 and HPAC cells were treated with Integrin β1 or Integrin β4 function blocking antibodies or control Mouse IgG1 in CAF-CM. NC media were used as a control. **g,** Cells were immuno-stained for p-NDRG1 T346 (white). Scale bar: 100 µm. **e-f,** Mean fluorescence intensities of p-NDRG1 signal were quantified in Fiji, plotted, and analysed using Mann-Whitney test. >150 cells were analysed per condition. Magenta lines represent medians. **h-i**, Src-family inhibitor Saracatinib (2 µM) or FAK inhibitor Defactinib (2 µM) were added to serum starved cells (SW1990, HPAC) 1h before addition of CAF-CM, the cells were lysed 3h later and immunoblotted for p-NDRG1 T346, NDRG1, p-FAK Y397, p-Src Y418, and tubulin. **j-k**, SW1990 and HPAC cells were treated with vehicle control (DMSO) or inhibitors targeting AKT (MK-2206 250 nM) or SGK1 (BLU6340 250 nM) and CAF-conditioned media was added for indicated times. Cells were lysed and probed for p-NDRG1, NDRG1, p-AKT S473 and vinculin.

First, we explored the kinetics of this phosphorylation in response to CAF-CM (Extended Data Fig. 2a-b) and observed upregulation of both total and phospho-NDRG1 in response to CAF-CM. We next aimed to determine the signalling events involved in phospho-NDRG1 up-regulation. Given our data showing high ECM secretion by the CAFs upon serum deprivation (Fig. 1a, Extended Data Fig. 1c-d), we hypothesized that ECM proteins might stimulate NDRG1 phosphorylation. Indeed, the addition of multiple soluble ECM proteins stimulated p-NDRG1 (Fig. 2c). Integrins are the major protein family sensing ECM proteins, and their engagement with ECM activates c-Src and FAK kinases^39^. Therefore, we assessed whether the application of integrin β1 or β4 -blocking antibodies or pharmacological inhibition of c-Src and FAK would lead to a decrease in p-NDRG1 (Fig. 2d). As hypothesized, blocking integrin β1 or β4 led to decreased p-NDRG1 (Fig. 2d). As hypothesized, blocking Integrin β1 or β4 led to decreased p-NDRG1 levels in the presence of CAF-CM (Fig. 2e-f), and inhibition of c-Src and FAK prevented NDRG1 phosphorylation upon addition of CAF-CM (Fig. 2h-i). Additionally, overexpression of Src and FAK kinases in the absence of ECM proteins or CAF-CM, also reduced the levels of DNA damage in PDAC cell lines treated with chemotherapy (Extended Data Fig. 2c-f). Thus, these data identify the integrin-Src-FAK signalling axis as an activator of ECM-induced NDRG1 phosphorylation and suggest that multiple ECM proteins can stimulate NDRG1 phosphorylation.

NDRG1 is phosphorylated by multiple kinases at its C-terminus, and the site that the RPPA detects (Thr346), can be phosphorylated by AKT (Protein kinase B) and SGK (Serum and Glucocorticoid induced kinases), which are closely related serine/threonine kinases belonging to the AGC family^40–42^. The C-terminus of NDRG1 contains more than 20 Ser and Thr residues, organised into three tandem repeats and the anti-pNDRG1 Thr346 antibody also recognizes the Thr356 and Thr366 phospho-sites; hence, assessment of specific NDRG1 phosphorylation sites is challenging. To investigate the role of phosphorylation on NDRG1 function and to identify the kinase responsible for NDRG1 phosphorylation in response to CAF-CM, we used the AKT inhibitor MK-2206 and a novel SGK1-3 selective inhibitor BLU6340 from Blueprint Medicines (data not shown for structure and syntheses in pre-print version). Inhibition of SGK1-3 abolished p-NDRG1 at Thr346 in the presence of CAF-CM while MK-2206 had no effect (Fig. 2j-k, Extended Data Fig. 2g-h), suggesting SGK1-3 as main kinases phosphorylating NDRG1 Thr346/356/366 in PDAC cell lines. We further noted that NDRG1 phosphorylation seemed to correlate with an increase in total NDRG1 levels over time (Fig. 2b-c, j-k, Extended Data Fig. 2a). Both SGK and NDRG1 have been reported to localize to the nucleus and cytoplasm^43–47^. Interestingly, SGK nuclear localization is dependent on its phosphorylation and activation and is predominantly nuclear during S phase^47^. To assess whether SGK inhibition might influence NDRG1 sub-cellular localization we performed IF analysis of cells treated with SGK inhibitor BLU6340. These data show that SGK inhibition results in a reduction of the nuclear NDRG1 pool (Extended Data Fig. 2i-k).

### NDRG1 expression correlates negatively with patient survival and efficacy of gemcitabine treatment

Because we observed that ECM proteins stimulate NDRG1 phosphorylation, we sought to investigate the role of NDRG1 in desmoplastic tumours. NDRG1 has been described as both a metastasis suppressor^31–33^ and as a tumour-promoting protein^34–37^. To reconcile these conflicting reports, we first investigated NDRG1 levels in human PDAC tissue microarray (TMA) PA805b (80 cases) with normal pancreas (20 cases) as a control. These data revealed that phospho- and total-NDRG1 were significantly elevated in PDAC tumours compared to normal tissue (Fig. 3a-c). 80% of normal tissues were negative for phospho-NDRG1 (Fig. 3b), while almost 50% of PDAC cases expressed phospho-NDRG1 (p=0.0354) (Fig. 3b). In contrast, 70% of all tissues stained positive for some level of total NDRG1, yet 82% of cancer cases exhibited moderate to strong (score 2 or 3) staining intensity, which was significantly higher than normal cases (p<0.001) (Fig. 3c).

**Fig. 3.**
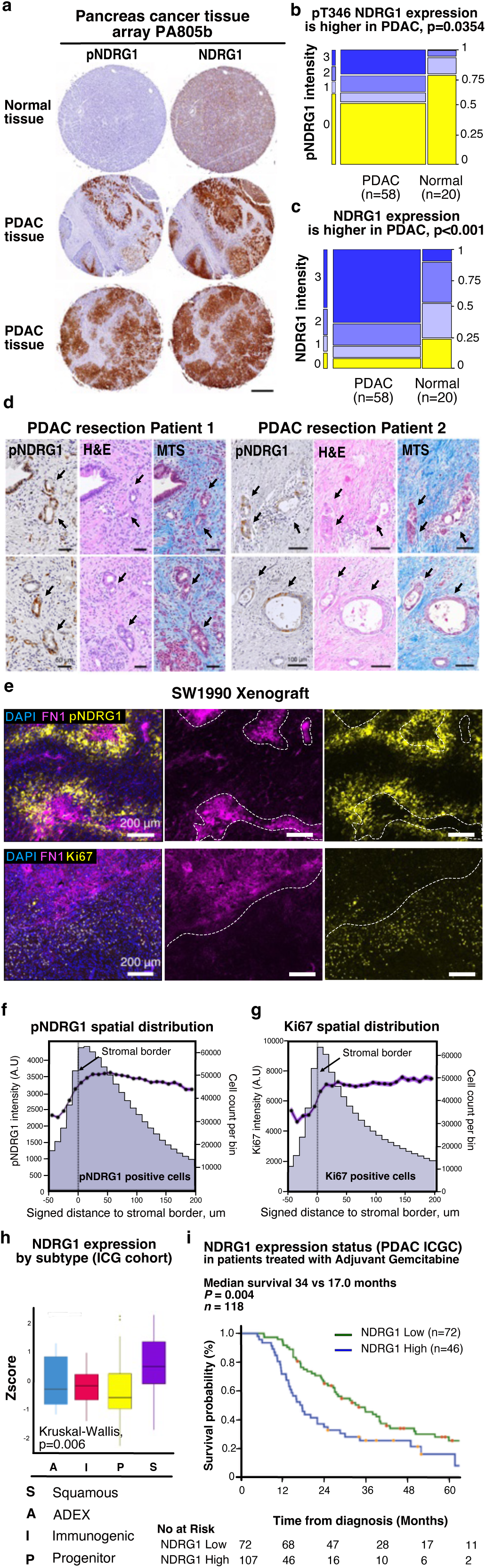
NDRG1 is highly expressed in PDAC and correlates with poor survival. **a,** Normal pancreas and PDAC tissue microarrays (PA805b, 20 normal, 58 tumours) were stained for phospho- and total NDRG1 expression, analysed for presence or absence of staining and graded by a pathologist. Representative images of IHC staining showing NDRG1 expression in normal vs. tumour tissue. Scale bar: 500 µm. PDAC. **b-c**, pNDRG1 and tNDRG1 expression/intensity from the tissue microarray from (a) was analysed by Fisher’s exact test. **d**, PDAC tissue resections of the 59 y.o. male patient treated with 6 cycles of gemcitabine/Abraxane (Patient 1) and of the 37 y.o. female patient treated with 6 cycles of FOLFIRINOX and Stereotactic body radiation therapy (SBRT) (Patient 2) were stained for phospho-NDRG1 expression. Representative images are shown. Two areas of interest per patient are shown. From left to right: panels show pNDRG1 positivity both in the nuclei and the cytoplasm of the neoplastic epithelium surrounded by the stroma. Middle panel - haematoxylin and eosin stain of the same tissue. Right panel - Masson’s Trichrome staining (MTS) highlighting the fibrotic stroma (in blue), cytoplasm of the neoplastic epithelium (red), and nuclei (dark red). Arrows point to neoplastic glands. Scale bars: 50µm (Patient 1) and 100µm (Patient 2). **e**, Representative gemcitabine-treated SW1990 xenograft tumour sections used for high-content spatial image analysis, stained for Fibronectin (FN1: magenta), p-NDRG1 (yellow) and Ki67 (yellow). White dotted line indicates stroma border. Scale bar: 200µm. **f-g,** pNDRG1 and Ki67 positive/high intensity cells are localized in close proximity to the stromal border. Spatial analysis of pNDRG1 and Ki67 signal intensity in cancer cells given their signed distance to the closest stromal border (Distance in µm is on x-axis). Graphs zoom on events at a -50 to 200µm distance to signed stromal border. Negative distance indicates cells located within stromal regions, while positive distances show cells located outside of these regions. Cells positive for pNDRG1 or Ki67 were grouped into bins (in violet) based on distance from signed stroma border and cell number per bin is plotted on the right y-axis, bin size is 10µm. Purple sensogram represents average maximum cellular signal intensity of pNDRG1 or Ki67- positive cells (black dots; Standard Error of the Mean [SEM] indicated as purple overlays) per bin and values are plotted on the left y-axis. **h**, Analysis of NDRG1 expression in the ICG cohort RNA-seq samples of nontreated bulk resected PDAC samples stratified by Squamous, Pancreatic Progenitor, Immunogenic and Aberrantly Differentiated Endocrine Exocrine (ADEX) subtypes. Kruskal-Wallis was used to assess significance. **i**, Disease-specific survival probability of patients treated with adjuvant gemcitabine therapy stratified based on NDRG1 mRNA expression status (high vs. low). Median DSS was calculated. Kaplan-Meier survival analysis and log-rank rest were used to analyze disease-specific survival.

We then investigated the status of *NDRG1* gene alterations and expression in patient samples from The Cancer Genome Atlas (TCGA)^48^. In this dataset, (referred to as PAAD in TCGA) the *NDRG1* locus was amplified in 11% of PDAC patients, and this locus was also amplified in multiple other solid tumours (Extended Data Fig. 3a). *NDRG1* alterations co-occurred with *KRAS* and *TP53* mutations in PDAC (data not shown), and patients with *NDRG1* amplifications showed significantly reduced overall survival compared to patients without *NDRG1* locus alterations (Extended Data Fig. 3b). In addition, high levels of phospho-NDRG1 in patient tissues were associated with shorter median overall survival (Extended Data Fig. 3c).

To evaluate the status of NDRG1 phosphorylation in human tissues in more detail we analysed PDAC tissue resected from patients treated with standard-of-care regiments, (gemcitabine/Abraxane or FOLFIRINOX and stereotactic body radiation therapy). We observed that each of the cancerous lesions positive for pNDRG1 was surrounded by fibrotic stroma as expected (Fig. 3d). Given that NDRG1 phosphorylation is induced by the ECM/integrin/FAK/SGK1 signalling pathway (Fig. 2), we hypothesized that NDRG1 phosphorylation within tumour tissue might be dependent on the proximity of the tumour cells to ECM. Human pancreatic adenocarcinomas do not have a defined ‘tumour edge’ but rather present as clusters of heterogeneous neoplastic glands (often comprising ≤10% of the resection specimen) dispersed in a fibrotic stroma. This heterogeneity limited quantitative assessment, and so we studied the biology of NDRG1 phosphorylation in relation to stromal abundance in carcinoma cells by utilizing gemcitabine-treated SW1990 mouse xenografts (Fig. 3e-g, Extended Data Fig. 3d). Using quantitative high-content spatial analysis, we identified stroma-dense areas of the tumour and calculated distances from the stromal border to phospho-NDRG1 and Ki67 positive cells. We observed that the areas of the tumour proximal to stroma border showed the highest presence of phospho-NDRG1 signal, and these areas were highly proliferative as defined by Ki67 positivity (Fig. 3e-g, Extended Data Fig. 3d-h).

Since molecular stratification of PDAC is important for guiding future treatment strategies, we were interested in whether NDRG1 expression is characteristic of a certain PDAC subtype. Analysis of NDRG1 mRNA expression in tumour samples from the International Cancer Genome Consortium (ICGC)^16^ showed NDRG1 to be more highly expressed in the Squamous subtype (Fig. 3h), known to be associated with the worst PDAC prognosis^49^. The analysis of 237 patients with PDAC who underwent surgery followed by adjuvant treatment with gemcitabine (ICGC) revealed that patients with high NDRG1 expression had worse survival compared to patients with low NDRG1 expression (median disease-specific survival [DSS] 15.9 months and 25 months, respectively) (Extended Data Fig. 3i). To determine whether NDRG1 expression might be indicative of clinical response to gemcitabine, patients who received gemcitabine in the adjuvant setting were stratified into high and low-NDRG1 expressing groups. Indeed, gemcitabine-treated patients with high NDRG1 expression had significantly shorter disease-specific survival compared to the low NDRG1 expressing group (median DSS 17 months and 34 months, respectively) (Fig. 3i). Overall, the analysis of the two pancreatic cancer patient cohorts TCGA and ICGC (187 and 237 patients respectively), together with patient samples from PA805b tissue microarray, all demonstrate that high NDRG1 expression is a poor prognostic factor.

### Loss of NDRG1 sensitizes cells to chemotherapy and impairs tumour growth

Because we observed increased phospho- and total NDRG1 expression in patient tumours and shorter survival in gemcitabine treated patients with high NDRG1 expression, we hypothesized that NDRG1 or its phosphorylation might be important for pancreatic tumour growth or therapy resistance. We first assessed whether SGK1 inhibition by BLU6340, or SGK1-3 knockdown, would reduce CAF-stimulated protection from DNA damaging chemotherapies (GEM, SN-38, 5-FU). We observed that either SGK inhibition or SGK1 knockdown increased DNA damage in CAF-CM treated PDAC cells (Fig. 4a and Extended Data Figure 4a-c). After observing higher levels of DNA damage in cells treated with the SGK1 inhibitor, we assessed whether knockdown of NDRG1 would similarly sensitize PDAC cells to chemotherapy and increase DNA damage. NDRG1 knockdown led to an increase in yH2AX foci and higher DNA damage in alkaline Comet assay in response to gemcitabine and SN-38 (Fig. 4b-d, Extended Data Fig. 4d), while having very little to no effect in response to 5-FU (Fig. 4c, Extended Data figure 4b-d). In addition, knockdown of NDRG1 increased yH2AX focus formation in untreated cells, and this effect became more pronounced following gemcitabine or SN-38 treatment. These findings suggest that NDRG1 knockdown increases basal levels of DNA damage, which is further exacerbated in the presence of DNA damaging agents (Fig. 4b, d).

**Fig. 4.**
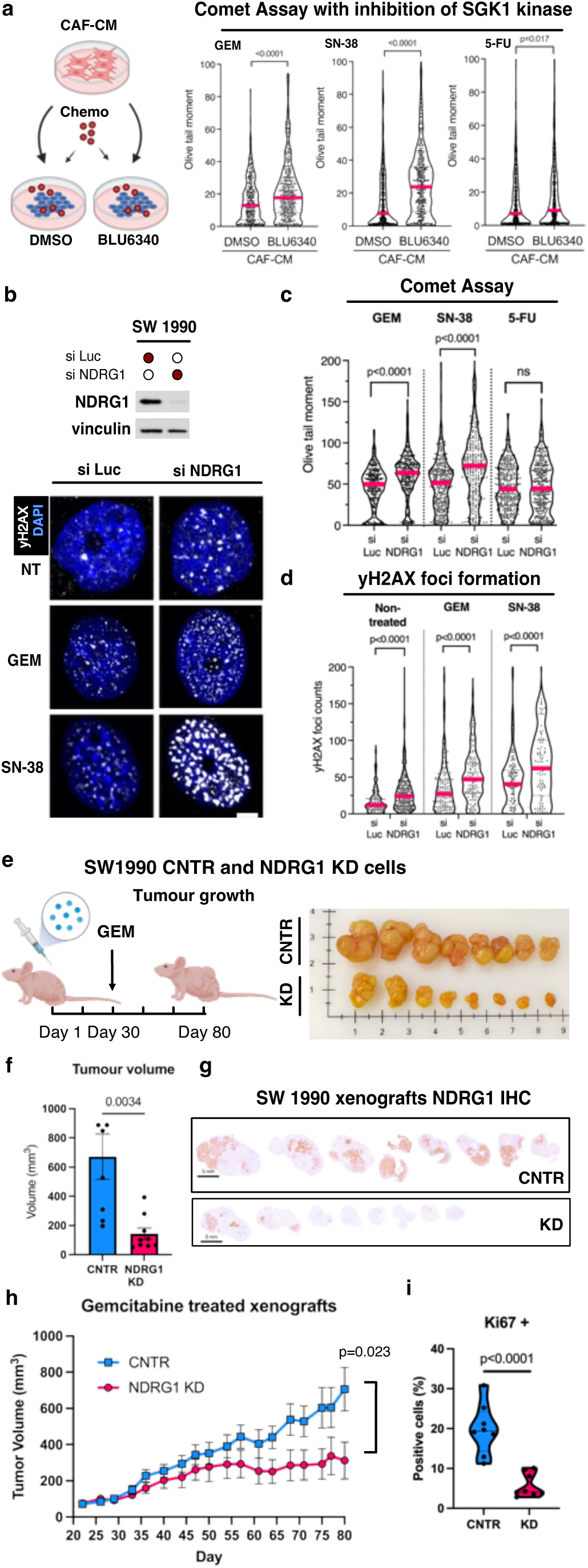
Loss of NDRG1 leads to higher DNA damage and eradicates CAF-CM protective effect to chemotherapy. **a,** SW1990 cells were treated with 1 µM gemcitabine (GEM), 0.5 µM SN-38, or 100 µM 5-FU in CAF-CM +/- 250 nM SGK1 inhibitor BLU6340. 24h later the cells were subjected to Comet assay. OTM values were analysed using two-tailed Mann-Whitney test, >150 cells per condition were analysed. Magenta lines represent median values. **b,** SW1990 cells were transfected with short interfering RNA (siRNA) against LUC or NDRG1 and probed for NDRG1 and vinculin. The cells were also treated with DMSO, 1 µM gemcitabine, or 0.5 µM SN-38, and immunostained for yH2AX (white) and DAPI (blue). Scale bar: 5µm. **c**, siLUC and siNDRG1 cells were treated with 1 µM gemcitabine, 0.5 µM SN-38, or 100 µM 5-FU in CAF-CM, and analysed by Comet assay. OTM values were plotted and two-tailed Mann-Whitney test was used for analysis. Magenta lines represent median values. **d**, yH2AX foci formation image quantification for (**b**) was performed using CellProfiler, foci numbers per single nucleus were plotted and two tailed Mann-Whitney test was used for analysis. Magenta lines represent median values. >100 cells were quantified per experiment. **e**, SW1990 cells expressing doxycycline-inducible shRNA against NDRG1 and CNTR cells expressing scrambled shRNA were injected into nude mice. NDRG1 knockdown was induced by feeding the mice doxycycline chow until the experimental endpoint. Tumours were allowed to grow until palpable and treated with 25mg/kg gemcitabine 2 times a week for 50 days. Tumours were harvested and imaged on Day 80. **f**, Tumour volumes were measured by calipers at the experimental end point and plotted. Student’s two-tailed T-test was used for statistics. **g,** SW1990 NDRG1 KD and CNTR IHC sections of the tumours were stained for NDRG1 expression. Scale bar: 5 mm. **h**, Growth curve of the tumours shown over time (NDRG1 KD: magenta, CNTR: blue). Student’s two-tailed T-test was used for statistics. **i,** Tumour sections were stained for pan-cytokeratin to define cancer cells, Ki67 to define proliferative cells, and the nuclei were counterstained by DAPI. Whole image analysis was performed in QuPath by applying appropriate thresholds to define pan-cytokeratin and Ki-67 positivity. Percent of the Ki67 positive cells in relation to pan-cytokeratin positive cells within each tumour is plotted. Student’s t-test was applied for significance. Error bars are SEM.

Next, to validate these data, we used CRISPR/Cas9 lentiviral delivery system to generate *NDRG1* knock-out (KO) cells (Extended Data Fig. 4e, h). *NDRG1* KO led to lower proliferation rates (Extended Data Fig. 4f, i), and sensitised the CAF-CM treated cells to gemcitabine and SN-38 (Extended Data Fig. 4g,j). These data suggest that NDRG1 is indeed necessary for CAF-stimulated chemoresistance.

Given that *NDRG1* KO cells proliferated more slowly, we anticipated that the SGK1 inhibitor would display a similar phenotype. However, to our surprise, the BLU6340 treated cells had identical proliferation dynamics to vehicle treated cells (Extended Data Fig. 4k-l). We next asked whether the BLU6340 treatment would sensitize cells to gemcitabine, and although we saw a shift in IC50 (Extended Data Fig. 4m), this was much more modest than what we observed with *NDRG1* KO lines (Extended Data Fig. 4g,j).

To understand these results, we assessed DNA damage in control (CNTR), *NDRG1* KO, or BLU6340 treated cells (SW1990 and AsPC) after gemcitabine or SN-38 treatment (Extended Data Fig. 4n). Although BLU6340 treated cells showed an increase in yH2AX foci compared to CNTR cells, *NDRG1* KO cells appeared to have an even higher yH2AX foci burden (Extended Data Fig. 4n).

Because most chemo-induced (and regular) DNA damage occurs in S phase, we wanted to assess whether SGK inhibition or NDRG1 deficiency would lead to changes in cell cycle dynamics that might explain our findings (Extended Data Fig. 4o-r). BLU6340-treated cells showed signs of replication stress judged by a lower number of cells in the S phase (Extended Data Fig. 4p, r), however *NDRG1* KO cells surprisingly had a higher S-phase cell number (Extended Data Fig. 4o, q). Given that NDRG1 loss leads to a slower proliferation rate (Extended Data Fig.4f, i) we hypothesized that lack of *NDRG1* might cause a specific S-phase defect. To assess this, we synchronized cells at the G1/S border and monitored cell cycle progression. We noticed that both BLU6340 treatment and NDRG1-deficiency led to a slower S-phase entry and an inability of cells to progress through S phase at the same pace as the CNTR cells (Extended Data Fig. 4o-r). However, BLU6340-treated cells were able to exit the S phase at the same time as CNTR cells judging by an increase in G2 and M phases of the cycle (10h time point for SW1990 and 8h for AsPC) (Extended Data Fig. 4p, r). In contrast, *NDRG1* KO cells were not able to complete the cell cycle at the same time as CNTR cells (Extended Data Fig. 4o, q). *NDRG1* KO cells showed prominent signs of accumulation in late S phase (8-12h time point for SW1990 and 5-9h for AsPC) and were not able to advance to G2/M as rapidly as CNTR cells. The evidence of replication stress in NDRG1 deficient cells, combined with their slower in S phase progression, might explain the heightened sensitivity of these cells to chemotherapies compared to BLU6340-treated WT cells.

Next, we wanted to assess the role of NDRG1 in tumour formation and chemoresistance in xenograft models. We injected SW1990 *NDRG1* WT and KO cells into nude mice and monitored tumour growth over time. While control cells (CNTR) formed distinct tumours (8/10), *NDRG1* KO cells were less tumourigenic (6/10) and significantly smaller in size (Extended Data Fig. 4s-u). We next asked whether NDRG1 knockdown (KD) leads to a better response to chemotherapy. We used SW1990 and AsPC CNTR and KD lines, injected them into nude mice, and once tumours were palpable, treated the mice with gemcitabine. We found that NDRG1 KD lines are more sensitive to gemcitabine, as shown by diminished tumour volume and reduction in Ki67+ positivity in comparison to CNTR cells (Fig. 4e-i, Extended Data Fig. 4v-z). These data collectively suggest that *NDRG1* loss leads to higher levels of DNA damage, sensitizes cells and tumours to DNA damaging agents, and impairs tumour formation.

### NDRG1 has a direct role in DNA replication and repair

Gemcitabine and irinotecan (SN-38) function by perturbing DNA replication, leading to replication stress through the stalling of replication forks and leading to the formation of double-strand breaks (DSBs)^2,50^. Since we observed that *NDRG1* KO removes the protective effect of CAF-CM, specifically against gemcitabine and SN-38-induced DNA damage and leads to an S-phase defect (Fig. 4), we hypothesized that NDRG1 might have a role in DNA replication, DNA repair, or in the protection of replication forks.

Previous work from the Cortez group had detected NDRG1 at active replication forks in HCT-116 cells by mass-spectrometry-coupled isolation of proteins on nascent DNA (iPOND)^51^. To determine whether NDRG1 has a role at replication forks in PDAC we also used the iPOND method^52^ (Fig. 5a). Interestingly, we detected both phosphorylated and total forms of NDRG1 not only at the sites of ongoing replication but also at stalled replication forks after treatment with Hydroxyurea (HU), whereas NDRG1 was absent from more mature chromatin (Fig. 5a). In line with a previous report^44^, we were able to detect the interaction of NDRG1 with PCNA, the DNA polymerase clamp that is a landmark of the replication fork, using Proximity-ligation assay (PLA). The number of PLA foci between NDRG1 and PCNA increased after replication fork stalling (Fig. 5b). To determine whether NDRG1 has a more direct role in DNA repair, we introduced siRNAs to knockdown NDRG1 in mouse embryonic stem cells that carry a homologous recombination (HR) reporter system (6×*Ter*-I-SceI-GFP) at the *Rosa26* locus, capable of measuring HR in response to site-specific replication fork stalling, or in response to a conventional site-specific DSB^53^ (Detailed in Extended Data Fig. 5a). The knockdown of NDRG1 reduced the efficiency of stalled fork HR by ∼50% but had no effect on DSB-induced HR (Extended Data Fig. 5b-c). Together, these data suggest that NDRG1 is present at replication forks and plays a specific role in stalled fork HR repair.

**Fig. 5.**
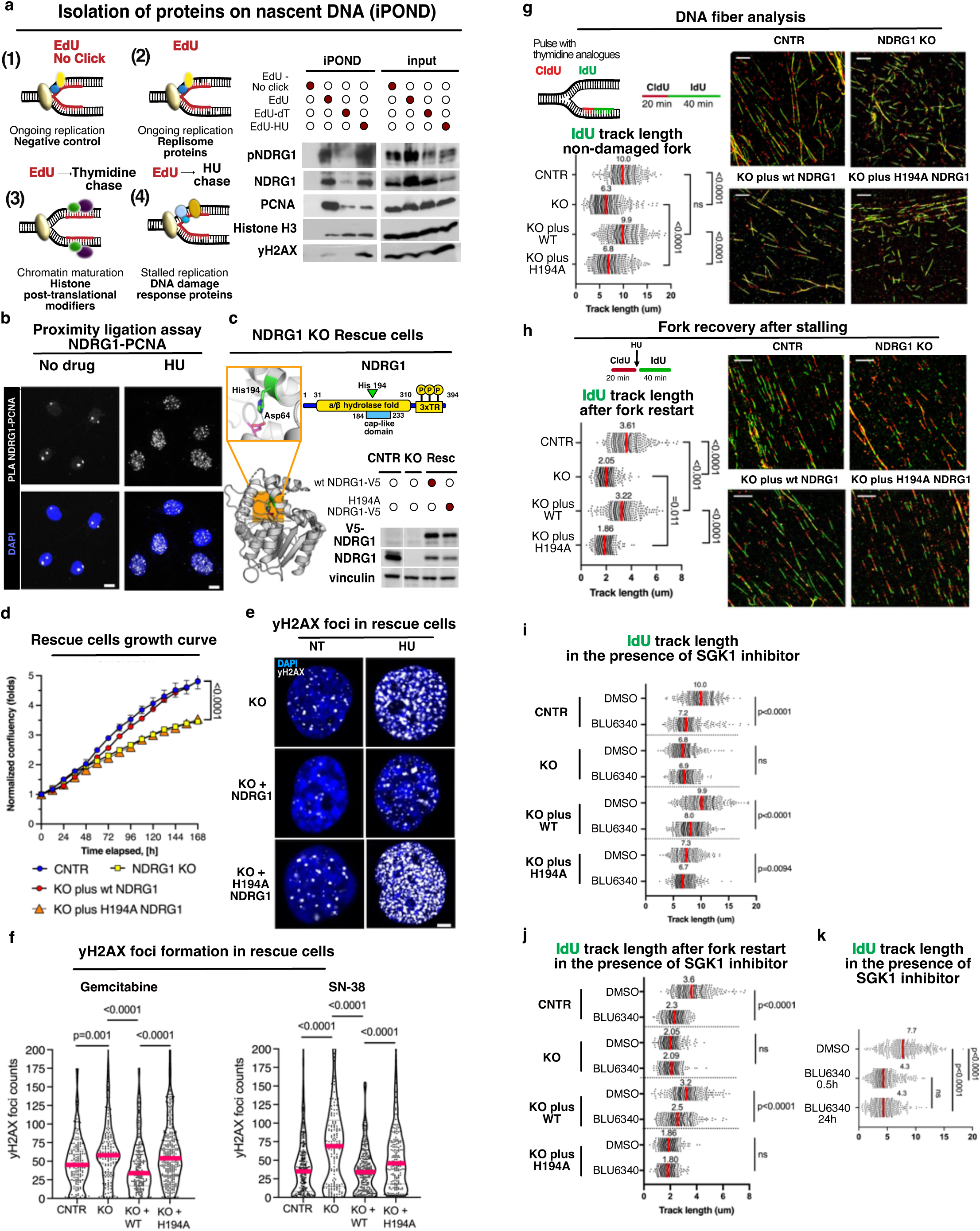
NDRG1 is found at the replication fork and regulates stalled fork recovery. **a**, Isolation of proteins on nascent DNA (iPOND). HPAC cells were used to assess proteins present at active replication fork (pulsed with 10µM EdU for 15min), during chromatin maturation (10 µM EdU 15min followed by 10 µM thymidine for 30min), and at stalled replication forks (10 µM EdU 15min followed by 4 mM hydroxyurea [HU]). EdU pulse without biotin azide present in Click-it cocktail was used as a negative control. iPOND fractions were run on SDS-PAGE and immunoblotted for pNDRG1 T346 and total NDRG1. PCNA, H3 and yH2AX were used as controls for various stages of replication fork progression/maturation. **b**, Interaction of PCNA and NDRG1 under conditions of active or stalled replication fork (HU 2 mM, 16h) was assessed by proximity ligation assay (PLA) in SW1990 cells. One dot indicates an interaction focus. Cells were fixed and immunostained according to the Duolink manufacturer’s protocol. Scale bar: 10 µm. **c,** Schematic of human NDRG1 where His194 (green) and the three phospho-residues in the C-term tandem repeats (TR) are marked. H194A mutation was introduced, and constructs expressing V5-tagged WT or H194A *NDRG1* variants were used for rescuing the *NDRG1* KO in SW1990 cells. Protein expression was probed by immunoblotting for V5 tag and NDRG1. **d**, Live cell proliferation analysis of CNTR (blue), *NDRG1* KO mix (yellow) and NDRG1 rescue cells (WT red, H194A orange) in 10% FBS. Error bars are SEM, Student’s t-test was used to establish significance. **e**, SW1990 *NDRG1* KO and rescue cells (WT and H194A) were plated on coverslips, treated with 2 mM HU for 24h, fixed and immunostained for yH2AX (white). Representative images of non-treated cells and HU-treated cells are shown, scale bar: 5µm. **f**, Quantification of yH2AX formation in SW1990 CNTR, *NDRG1* KO, and rescue (WT, H194A) cells after 1µM gemcitabine or 0.5µM SN-38 treatment (24h). yH2AX foci quantification was performed using CellProfiler, foci numbers per single nucleus were plotted and two tailed Mann-Whitney test was applied to establish significance. Magenta lines represent median values. >150 cells were quantified per experiment. **g**, Replication fork dynamics were analysed by DNA Fiber Assay. CNTR, *NDRG1* KO and *NDRG1* rescue cells (WT and H194A) were labelled with CldU followed by IdU as indicated. Median IdU track lengths (µm) of >250 double-labelled fibers are shown in magenta. Mann–Whitney test was applied to test significance. Representative images are shown, scale bar: 10µm. **h**, To assess fork recovery after stalling, CNTR, *NDRG1* KO and *NDRG1* rescue cells (WT and H194A) were labelled with CldU, treated with 2 mM HU for 1h to stall the replication fork, washed and IdU pulsed as indicated. Median IdU track lengths (µm) of at least 240 double-labelled DNA fibers, indicating fork restart are indicated in magenta. Representative images are shown, scale bar: 5µm. **i,** To analyse the effect of SGK1 inhibition on replication fork dynamics, SW1990 CNTR, *NDRG1* KO mix and *NDRG1* rescue cells (WT and H194A) were labelled with CldU and IdU as indicated in the presence of SGK1 inhibitor BLU6340 (250 nM, for 24h). IdU tracks were measured (µm) and medians indicated in magenta. Mann–Whitney test was used for significance. **j**, To assess how SGK1 inhibition influences fork re-start after stalling, cells were treated with 2mM HU as indicated in the presence of SGK1 inhibitor BLU6340 (250 nM), IdU track were measured and plotted. Scale bar: 5 µm. Medians indicated in magenta. Mann–Whitney test was used for significance. **k,** To analyse the effect of short-term vs. long-term SGK1 inhibition on replication fork dynamics SW1990 cells were pulsed with CldU and IdU as described in (**g**) and treated with either vehicle control (DMSO), or 250nM BLU6340 for 24h or pre-treated with BLU6340 for only 0.5h, after which the DNA fibers were analysed. Mann–Whitney test was applied to test significance, and magenta line indicates median.

NDRG1 is considered to be a potentially inactive member of the α/β hydrolase enzyme family because it lacks the canonical α/β hydrolase catalytic triad residues^54^. However, the crystal structure of the NDRG1 hydrolase fold (aa31-319) revealed putative enzyme active site residues, Asp64 and His194, located near the position of the canonical α/β hydrolase catalytic Ser-His-Asp triad between the α/β hydrolase core and the cap domain^55^. To examine whether His194 is important for the NDRG1-mediated reduction of DNA damage, we generated the H194A mutation and re-expressed either the WT *NDRG1* or the H194A *NDRG1* in the *NDRG1*-null background (Fig. 5c). While the proliferation rate of the cells re-expressing WT *NDRG1* was similar to control cells, strikingly, the proliferation rate of the H194A *NDRG1* cells phenocopied the *NDRG1* KO although the protein was expressed at similar levels as the WT add-back (Fig. 5c, d). Furthermore, when we analysed yH2AX levels in the *NDRG1* KO cells or in cells re-expressing the WT or the H194A mutant after treatment with SN-38, gemcitabine, or HU, we observed that WT *NDRG1* rescued the DNA damage effect, whereas the H194A mutant failed to do so even in the absence of any DNA damaging agent (Fig. 5e-f, Extended Data Fig. 5d). These data suggest that His194 is an important residue for NDRG1’s role in DNA repair.

Since *NDRG1* KO and H194A *NDRG1*-expressing cells grew more slowly than the wild type cells, we monitored replication fork progression using the DNA fiber spreading technique (Fig. 5g, Extended Data Fig. 5e)^56^. SW1990 and AsPC CNTR, NDRG1 KO, and KO add-back cells (WT or H194A), were pulsed successively with two thymidine analogues (CldU and IdU), and the IdU-labeled DNA fiber tracks were measured. NDRG1 knockout led to shorter IdU fiber track length further confirming that NDRG1 loss causes replication stress. While re-expression of WT *NDRG1* reverted the fiber track lengths back to CNTR cells length, H194A *NDRG1* re-expressing cells were unable to rescue the replication fork speed back to control level. This result suggests that H194A *NDRG1* cells, similarly to KO cells, undergo perturbed replication (Fig. 5g, Extended Data Fig. 5e). To further investigate the role of NDRG1 in stalled fork repair, we stalled the replication fork after the CldU pulse by adding HU for 1h, followed by release from HU into IdU pulse. *NDRG1* KO cells had shorter IdU tracks than CNTR cells, suggesting that replication restart after HU was impaired. This defect was fully rescued by the re-expression of WT *NDRG1*. In contrast, re-expression of H194A *NDRG1* failed to restore normal fork restart kinetics compared to control cells (Fig. 5h, Extended Data Fig. 5e). Taken together, these data indicate that His194 is a critical residue for NDRG1 function in DNA replication and DNA repair.

To investigate whether inhibition of SGK results in slower fork progression, we treated the CNTR, *NDRG1* KO, and the add-back (WT, H194A) cells with the SGK inhibitor BLU6340, and analysed normal and stalled replication fork dynamics (Fig. 5i-k). Indeed, consistent with our previous observation that SGK inhibition leads to higher levels of DNA damage after gemcitabine and SN-38 treatment, and delayed S phase entry (Fig. 4), IdU tracks were shorter in the presence of BLU6340, mimicking the phenotype of *NDRG1* KO cells (Fig. 5g-h, Extended Data Fig. 5e). BLU6340 had similar effects on fiber length in KO cells rescued with WT *NDRG1* but failed to reduce fiber lengths further in the H194A add-back cells. SGK inhibition also impaired replication fork recovery in CNTR cells and WT *NDRG1*-rescued cells but had very little effect on *NDRG1* KO cells or H194A cells (Fig. 5i-j). These findings reveal an epistatic relationship between SGK1 inhibition and NDRG1 mutation in replication stress phenotypes. This, in turn, suggests that NDRG1 is the critical target of SGK1 phosphorylation in the suppression of replication stress. To rule out long-term effects of SGK1 inhibition on potential replication fork perturbations and total NDRG1 levels, we analysed how short- (30 min) or long-term SGK1 inhibition (24 h) influences fork progression. We found that both short- and long-term BLU6340 treatment results in slower fork progression (Fig. 5k, Extended Data Fig. 5f), underscoring the importance of SGK-mediated NDRG1 phosphorylation for normal replication.

Finally, because we observed that CAF-CM stimulates NDRG1 phosphorylation via SGK1, we wanted to analyse the effects of CAF-CM and SGK inhibition on fork dynamics, and whether any such effect occur via NDRG1. We analysed the effect of CAF-CM on non-damaged and stalled fork dynamics in the presence or absence of NDRG1, or with and without BLU6340 (Extended Data Fig. 5g-h). Consistent with our findings, fiber spreading analysis revealed that CAF-CM indeed increases replication fork speed and stimulates recovery from fork stalling. In addition, the CAF-CM effect on replication fork dynamics is dependent on NDRG1 and on NDRG1 phosphorylation by SGK1.

These data suggest that NDRG1 and its phosphorylation by SGK1 are important for normal fork progression and stalled fork repair. We further show that mutation of NDRG1 His194 to Ala, phenocopies the loss of *NDRG1*, rendering NDRG1 unable to maintain replication fork homeostasis.

### NDRG1 prevents R-loop accumulation

Replication fork slowing and stalling can be caused by many factors. To determine whether the replication fork defects caused by NDRG1 loss reflect overall slowing of global replication, or increased pausing at replication fork barriers, we analysed sister fork symmetry^57^. If there were an overall slowing of replication, we would anticipate both sister forks to slow at a similar rate, maintaining symmetry. However, if increased fork pausing were the major mechanism of slowed replication, we would expect to observe an asymmetric pattern of sister replication tracts (Fig. 6a). Using the DNA fiber assay, we found that NDRG1-depleted or BLU6340-treated cells show substantially increased asymmetry of sister replication tracts, compared to control cells (Fig. 6a). Our data therefore suggest that NDRG1 deficiency increases the frequency of replication fork stalling events.

**Fig. 6.**
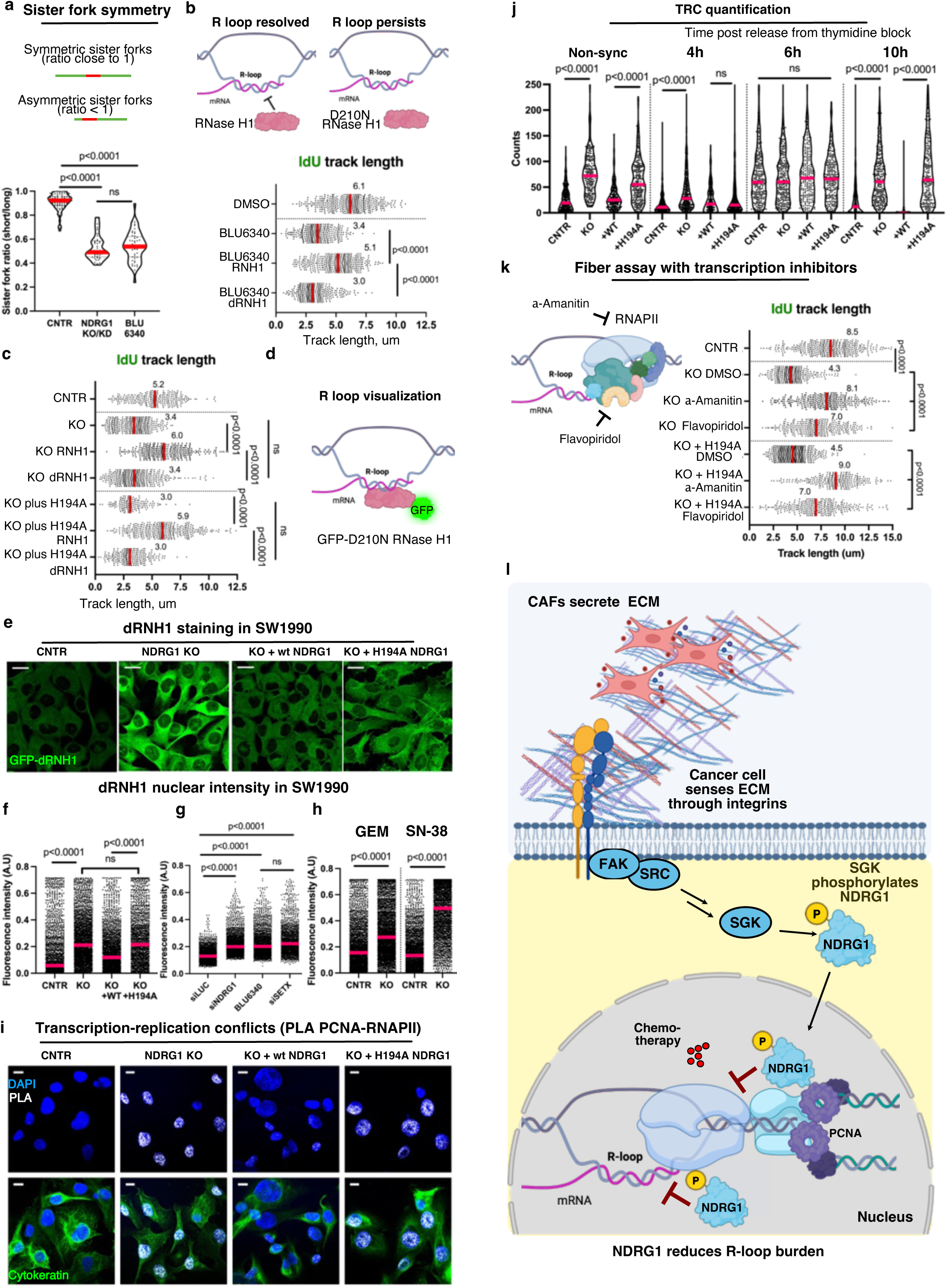
NDRG1 is involved in resolving R-loops. **a,** Schematic for the sister fork symmetry analysis. Asymmetric sister forks show a ratio of a shorter IdU track to longer track <1, while symmetric forks show ratio ≈ 1. SW1990 CNTR, NDRG1 deficient and cells treated with 250nM BLU6340 were analysed for sister fork symmetry. 89 bidirectional forks were analysed for CNTR cells, 44 for NDRG1 deficient and 49 for BLU6340-treated cells, magenta lines represent median, Mann-Whitney test was used for significance. **b,** Schematic of the RNase H1 constructs used in the assays, WT-RNase H1 removes R-loops whereas the catalytically dead D210N RNase H1 is unable to do so. Replication fork progression was analysed using DNA fiber assays. SW1990 cells were treated with vehicle control (DMSO) or 250nM BLU6340 and co-transfected with constructs encoding WT-RNase H1 (RNH1) or the catalytically dead (dRNH1). Cells were pulsed with IdU (20min) followed by CldU (40min). Median IdU track lengths (µm) of >300 double-labelled fibers are shown in magenta. Mann–Whitney test was applied to test significance. **c,** Fork dynamics were analysed using DNA fiber assays. SW1990 CNTR, *NDRG1* KO, and H194A addback cells were used and transfected with either WT-*RNH1*, or dead *RNH1* (dRNH1). Median IdU track lengths (µm) of >150 double-labelled fibers are shown in magenta. Mann–Whitney test was applied to test significance. **d,** Schematic of the recombinant RNase H1 construct used for R-loop visualisation. The bacterial purified GFP-labelled catalytic dead (D210N) RNase H1 was used to probe for RNA-DNA hybrids in fixed cells. **e**, Confocal images of SW1990 cells stained with GFP-dRNH1 (green) in CNTR, NDRG1 KO and rescue (WT, H194A) SW1990 cells. Scale bar: 20µm. **f,** Analysis of the fluorescence intensity of nuclear GFP-dRNase H1 in SW1990 cells from (**e**). At least 1500 nuclei were analysed per condition, magenta lines represent median, Mann-Whitney test was used for significance. **g,** GFP-dRNH1 nuclear intensity was analysed in SW1990 CNTR cells, cells with NDRG1 knockdown (siNDRG1), or after BLU6340 250nM 24h. Knockdown of Senataxin (siSETX) was used a positive control to induce R-loops. At least 1600 nuclei were analysed per condition, magenta lines represent median, Mann-Whitney test was used for significance. **h,** To assess whether *NDRG1* KO increases R-loop content in the presence of chemotherapies SW1990 CNTR and KO cells were treated with 0.5µM gemcitabine (GEM) or 0.5 µM SN-38 for 24h and analysed for RNA-DNA hybrid formation by GFP-dRNH1 staining. At least 2100 nuclei were analysed per condition, magenta lines represent median, Mann-Whitney test was used for significance. **i,** Confocal images of the transcription-replication conflicts (TRC) in SW1990 cells. Proximity ligation assay (PLA) between PCNA and RNA PolII (RNAPII) was performed in CNTR, *NDRG1* KO and rescue (WT, H194A) cells. PLA foci between PCNA and RNAPII are shown in white. Cytokeratin (green) was used to visualize cell shape, Scale bar: 10µm. **j,** SW1990 CNTR, *NDRG1* KO, and rescue (WT, H194A) cells were synchronized by double thymidine block at G1/S border and TRC formation was analysed at different time-points post dT release. TRC accumulation was also analysed in non-synchronized cells. TRCs were assessed by PLAs between PCNA and RNA PollI, imaged and TRC foci counts analysed by CellProfiler. At least 210 nuclei per timepoint were analysed, magenta lines represent median, Mann-Whitney test was used for significance. At 6h all the lines had significant TRC load which were resolved in the CNTR and WT add-back cells at 10h timepoint. **k,** DNA fiber assay in SW1990 CNTR, *NDRG1* KO and H194A add-back cells in the presence of transcription inhibitors α-amanitin (10mg/ml) and flavopiridol (10µM). Transcription inhibitors reduce TRC conflicts, thus preventing R-loops accumulation and rescuing replication fork dynamics in NDRG1-deficient or H194A NDRG1 expressing cells. Cells were pulsed with CldU (20min) and IdU (40min). Median IdU track lengths (µm) of >275 double-labelled fibers are shown in magenta. Mann–Whitney test was applied to test significance. **l,** Schematic of the proposed mechanism by which ECM proteins stimulate DNA repair and reduce R-loop formation via integrin-FAK-SRC-SGK1-NDRG1 pathway.

One of the major barriers to normal replication fork progression and a recognized source of replication stress is the formation of RNA-DNA hybrids (‘R-loops’) during transcription^58–60^. To investigate whether R-loops might contribute to the replication stress and fork stalling observed in NDRG1 null cells, BLU6340-treated cells, or *NDRG1* KO cells re-expressing *NDRG1* H194A, we expressed either wild-type *RNase H1* (an endonuclease able to eliminate RNA-DNA hybrids), or catalytically inactive *RNase H1* (D210N or dRNH1), and measured IdU track lengths. We found that WT RNase H1, but not dRNH1, was able to restore fiber length in the *NDRG1*-perturbed cells (Fig. 6b-c, Extended Data Fig. 6a-b). These data suggest that R-loops are a major contributor to the fork stalling observed in *NDRG1*-perturbed cells.

To determine whether the abundance of R-loops was affected by NDRG1 status, we used a recombinant catalytically dead GFP-tagged *RNase H1* (GFP-dRNH1) as a probe for R-loop abundance^61^. We found that NDRG1-deficient cells, cells treated with BLU6340, or cells expressing *NDRG1* H194A have higher levels of nuclear RNA-DNA hybrids compared to control cells (Fig. 6d-g, Extended Data Fig. 6c, e-g). Similar data were obtained using the S9.6 antibody that also recognizes RNA-DNA hybrids (Extended Data Fig.6i-j).

TOP1 inhibitors cause an increase in R-loops, reflecting a requirement for TOP1 to relax DNA supercoils generated during transcription ^59^. We wanted to assess to what extent the chemotherapies we were using, SN-38 (a TOP1 inhibitor) and gemcitabine (a non-TOP1 inhibitor), would increase R-loops in PDAC cells. We did observe a significant increase in R-loops in both chemo-treated cells (Extended Data Fig. 6d) and *NDRG1* loss further exacerbated R-loop burden compared to the CNTR cells (Fig. 6h, Extended Data Fig. 6h).

R-loop-induced fork stalling is a major feature of transcription-replication conflicts (TRCs) and R-loop-induced DNA damage in S/G2 phase^62,63^. Collisions between PCNA and RNAPII can be assessed by proximity ligation assay (PLA). Indeed, we observed a higher number of TRCs in *NDRG1* KO cells, and in *NDRG1* KO cells re-expressing *NDRG1* H194A (Fig 6i-j, Extended Data Fig. 6k-l). Given the impaired S phase in NDRG1-defective cells, we investigated the dynamics of TRC formation and resolution during cell cycle progression (Fig 6j, Extended Data Fig. 6l). We found similar levels of TRCs in S phase in all the cells studied (CNTR, *NDRG1* KO, and *NDRG1* KO with WT addback or *NDRG1* H194A addback). However, only the CNTR and WT *NDRG1* add-back cells were able to resolve these TRCs as the S phase progressed. Cells with defective NDRG1 function revealed sustained levels of TRCs during S phase progression. The persistence of TRCs, and the consequent defects in replication fork progression may explain the prolonged S phase we observed upon NDRG1 loss.

If the failure to resolve TRCs were a major cause of replication stress in NDRG1-defective cells, blocking transcription might be predicted to rescue normal replication speed in *NDRG1* KO cells. Indeed, we observed that general transcription inhibitors, α-amanitin and flavopiridol, were able to revert fiber lengths of NDRG1 mutants closer to WT length (Fig. 6k, Extended Data Fig. 6m). α-amantin had a more pronounced effect than flavopiridol, likely because flavopiridol, a CDK9 inhibitor (which also targets other CDKs), does not fully inhibit transcription, whereas α-amantin is a highly selective inhibitor of POLII and POLIII^64^.

In summary, this study reveals an unexpected connection between CAF-secreted ECM proteins, signalling through the Integrin/Src/FAK/SGK1 axis, and NDRG1-mediated suppression of replication stress. We further show that NDRG1 acts at the replication fork, participates in repair of stalled replication forks and reduces the R-loop burden during S phase (Fig. 6l).

## Discussion

Our study offers novel mechanistic insight into how CAF-secreted extracellular matrix proteins induce chemoresistance via the reduction of DNA damage in pancreatic ductal adenocarcinoma (Fig.6l). We show that lipid and serum starvation leads to enhanced ECM secretion by CAFs, which stimulates NDRG1 phosphorylation by SGK in response to integrin/Src/FAK-signalling (Fig. 1-2). Our analysis of two pancreatic cancer patient cohorts identified elevated NDRG1 expression level as a prognostic factor for poor overall survival outcome and chemotherapeutic response (Fig. 3). We further show that phosphorylated NDRG1 is present at active and stalled replication forks and prevents replication forks from stalling by reducing the R-loop burden. These activities likely explain how NDRG1 mediates resistance to chemotherapies (Fig. 4, Fig. 5, Fig. 6). Since tumours treated with chemotherapies are reliant on replication fork homeostasis, increased genomic instability caused by *NDRG1* loss leads to overall lower tumour fitness and increased sensitivity towards replication fork stalling agents.

Elevated ECM deposition, high stromal density and altered mechanical properties of the TME cause resistance to multiple chemotherapies in the cancers of the breast, pancreas, lung and ovaries^5,6,65–76^. To our knowledge, this study is the first one to clearly link the ECM environment to DNA repair at a mechanistic level. NDRG1’s role in cancer has been elusive since NDRG1 has been described as a metastasis suppressor^31–33^ and as a tumour-promoting protein^34–37^. Some reports that show NDRG1 expression in PDAC lead to a more favourable outcome^77^. However, our analysis of the TCGA and ICGC data shows NDRG1 to be highly amplified and overexpressed in many solid tumours, particularly in PDAC. We hypothesize that the discrepant conclusions reached regarding NDRG1’s role in tumorigenesis might result from studies of different cancer lineages, and further studies are required to establish the dualistic nature of NDRG1. For example, NDRG1 is highly expressed in the aggressive squamous/basal subtype of PDAC, a subtype characterized by high replication stress signature^78^, which might explain differential dependence on NDRG1 in different tumour contexts.

High NDRG1 expression has been previously linked to resistance to anti-cancer therapies^79^, such as alkylating chemotherapy temozolomide in glioma through stabilization of O^6^-methylguanine DNA methyl transferase (MGMT)^38^. Also, in line with our findings, that study found that phosphorylation of NDRG1 by SGK1 was required for NDRG1 interaction with MGMT and other DNA repair machinery proteins^38^. In addition, NDRG1 was shown to drive radiotherapy resistance in colorectal cancer^80^, and doxorubicin resistance in hepatocellular carcinoma^81^. These observations are further supported by our analysis of the largest up-to-date pancreatic cancer patient cohort, where patients with high *NDRG1* expression had worse disease-specific survival after adjuvant gemcitabine treatment (17 months vs 34 months respectively) (Fig. 3). Additional research is needed to differentiate whether these effects are due to more aggressive behaviour in NDRG1-high tumours, NDRG1-induced resistance to therapy, or both.

Genomic instability can lead to a variety of complications, from cancers to neurological disorders and mutations in DNA damage response proteins have been linked to various neurodegenerative diseases^82^. Mutations in the hydrolase domain of NDRG1 are known to cause Charcot-Marie-Tooth disease type 4D (CMT4D) neuropathy. Interestingly, a genome-wide siRNA screen in HeLa and U2OS cells revealed higher levels of yH2AX foci signal upon the knockdown of several genes causing Charcot-Marie-Tooth disease, including NDRG1^83^. This connection opens new avenues of research on CMT predisposition genes and their role in DNA repair and cancer. It is possible that CMTD4D mutations might impair NDRG1 DNA repair functions or its interaction with other DNA repair proteins, causing elevated levels of DNA damage and potentially contributing to this neuropathic disorder.

Besides causing replication stress and higher levels of DNA damage^63^, dysregulated formation of R-loops is also associated with several neurological disorders^84^. R-loop burden can be controlled at many levels. Both their formation and resolution involve a variety of molecular players, including RNA-dependent ATPases (SETX, FANCM, AQR, DDX19, DDX5, BLM and others) that resolve R-loops by using a DNA-RNA unwinding activity^59^. Interestingly, knockdown of *NDRG1* or BLU6340 treatment led to a similar R-loop burden as the one caused by the *SETX* knockdown (Fig. 6g, Extended Data Fig. 6g). Since we find that *NDRG1* KO and H194A *NDRG1* expressing cells are unable to resolve TRCs, we postulate that NDRG1 normally prevents R-loop accumulation.

Despite the reports that NDRG1 is devoid of catalytic activity^54^, a structural study allowed speculating about possible catalytic activity of NDRG1^55^. The substrate, if any, for NDRG1’s putative activity is yet to be uncovered. However, our study shows that the H194A NDRG1 mutant failed to rescue multiple DNA damage phenotypes, whereas WT NDRG1 did (Fig. 5-6). In addition, phosphorylation of NDRG1 by SGK1 appears to also be important for the proper function of NDRG1 in replication fork homeostasis. Since the crystal structure of the NDRG1 variant (aa31-319) did not contain the extended, flexible C-terminus (including the SGK1 sites at Thr346, Thr356, Thr366), we can only speculate that in the absence of the C-terminal phosphorylation, NDRG1 might either not attain full function, not localize to the nucleus properly or as reported previously, might be unable to bind other interactors necessary for its DNA repair functions^38^. Resolving these structure-function questions regarding NDRG1 will require future studies.

Overall, our data identified mechanisms of NDRG1 function at replication forks and encourages the development of novel treatment strategies for PDAC patients with high NDRG1 expression. This could be achieved by targeting DNA repair functions of NDRG1 in combination with other DNA damaging agents, fork stalling agents, and targeted therapies against molecular players involved in R-loop associated DNA damage.

## Materials and methods

### Chemicals

Biochemicals and enzymes were of analytic grade and were purchased from commercial suppliers. Gemcitabine (S1714), activated form of Irinotecan SN-38 (S4908), 5-FU (S1209), Oxaliplatin (S1224), MK-2206 (S1078), Saracatinib (S1006), Defactinib (S7654) were from Selleckchem. SYBR™ Gold Nucleic Acid Gel Stain (S11494) was from Ivitrogen. 5-chloro-2′-deoxyuridine (CldU) (C6891), 5-iodo-2′-deoxyuridine (IdU) (I7125), DAPI (D9542), hydroxyurea (H8627), thymidine (T1895), and 5-ethynyl-2′-deoxyuridine (EdU) (900584) were from Sigma. Restriction enzymes were from NEB.

Small molecule SGK1 inhibitor BLU6340 was made by Blueprint Medicines, Cambridge, MA, USA. The cell effects of BLU6340 were determined in MDA-MD-436 cells (ATCC:HTB-130). The cells were plated in 384 well plates at a density of 20,000 cells per well in RPMI-1640 media supplemented with 10% heat inactivated FBS. 24h post-plating, the cells were treated for 1 hour at 37°C with either DMSO or BLU6340 in a 10 point, 3-fold dose response with a top concentration of 10µM. The final DMSO concentration for all wells was 0.5%. After 1h compound treatment, the cells were lysed in lysis buffer #4 (CisBio) and both pNDRG1 (which is used as a read-out for SGK activity) and total NDRG1 were measured by a custom assay developed by Meso Scale Discovery. The average IC50 from 5 independent assays measuring BLU6340 mediated inhibition of pNDRG1 was 33.1nM with a STDEV of 20.4nM.

### Isolation and culture of primary human cancer-associated fibroblasts

Human pancreatic cancer-associated fibroblasts referred as CAF Patient 1 and CAF Patient 2 lines were a gift from Dr. Chris Halbrook. Isolation of human cancer-associated fibroblasts was performed as previously described^85^. The IRB UCI 08-70 was used to establish the CAF lines, and informed consent was obtained. Fresh surgical tissue samples from PDAC patients were minced into 1 mm^3^ pieces using a sterile scalpel. Several tissue pieces were placed in the middle of a 6-well plate and covered in 800 µl isolation medium (DMEM/F12, 20% FBS, GlutaMAX, HEPES, Penicillin-Streptomycin). Plates were placed into an incubator and medium was replaced after 24 hours. For subsequent culture, FBS-reduced growth medium (DMEM/F12, 10% FBS, GlutaMAX, HEPES, Penicillin-Streptomycin) was used. Once fibroblast outgrowth was observed, tissue pieces were moved within the plate using a sterile pipette tip to achieve an even distribution of cells. Cells were trypsinized when confluent and transferred to 10 cm dishes for expansion and freezing.

### Cell culture and conditioned media preparation

Pancreatic adenocarcinoma cell lines HPAC (CRL-2119), SW1990 (CRL-2172), BxPC-3 (CRL-1687) were purchased from ATCC. SUIT-2, AsPC, and PA-TU-8889T cell lines were a kind gift from Dr. Nada Kalaany. All cell lines were tested *Mycoplasma* negative by using the MycoAlert Detection Kit (Lonza) and STR authenticated. Immortalized human pancreatic stellate cells, the most relevant cell model to study the role of pancreatic cancer–associated stromal fibroblasts, referred to as CAFs in the study, were a gift from Dr. Rosa Hwang^86^. Pancreatic cancer cell lines and pancreatic CAFs were routinely maintained in DMEM supplemented with 10% FBS, 50 IU/ml penicillin, and 50 μg/mL streptomycin. In order to obtain conditioned media from CAFs, CAFs were plated onto 15cm plates, and when reached 90% confluency the media was changed from 10% FBS to 0% FBS. 3-4 days later, the media was harvested, spun down to remove cell debris, and sterile filtered through 0.22 µm filter. The media was either snap frozen in LN or used fresh. When indicated, pancreatic cancer cell lines were incubated in non-conditioned media (NC) (2% FBS), or in the conditioned media (CAF-CM 2% FBS) for the course of the experiment. Fractionation of the CAF-CM was performed via centrifugation using the 3-kDa cutoff filter (Amicon Ultra MWCO 3kDa).

### TCA precipitation and Western Blotting

Media from CAFs was mixed with 100% (w/v) TCA (trichloroacetic acid with the final concentration of the acid 20%, incubated on ice for 2h, spun at 14000 rpm for 10 min, pellets were washed with a solution of ice cold 0.01 M HCl / 90% acetone. Later the pellets were boiled in 2× SDS-loading buffer and separated by SDS-PAGE. The pancreatic cancer cells were lysed in RIPA buffer (BP-115, Boston Bioproducts), complete protease inhibitor cocktail tablet (Roche), kept on ice for 10 minutes, sonicated, and centrifuged at 14,000 × g for 10 minutes at 4°C. Proteins (20–50 μg per sample) were separated by electrophoresis on 4%–12% pre-cast Tris-Glycine gels (Invitrogen) and transferred to nitrocellulose membranes. The membranes were incubated with the following antibodies: Collagen 1 (Abcam #ab34710), Fibronectin 1 (Abcam #ab6328, Invitrogen #MA5-11981), pan-Laminin (Dako #Z-0097), EGFR (#sc-03G, Santa Cruz), GAPDH (#5174S, CST), phospho-NDRG1 T346 (#5482S, CST), NDRG1 (#9485S, CST), phospho-FAK Y397 (#3283S, CST), phospho-Src Y418 (#44-660G, Invitrogen), phospho-AKT S473 (9271, CST), V5-tag (R96025, Invitrogen), PCNA (#2586S, CST), Histone H3 (#4499S, CST), pH2AX S129 (yH2AX) (#9718S, CST and 05-636 Millipore Sigma), α-tubulin (B-5-1-2) (T5168, Sigma-Aldrich), β-actin (A5316, Sigma-Aldrich), vinculin (#13901, CST), integrin ⍺5 (#4705, CST), integrin ⍺V (#4711, CST), integrin β4 (#14803, CST, Abcam #ab110167), integrin ⍺4 (#8440, CST), integrin β1 (#9699, CST, BD clone 9EG7 #553715), integrin β1 active (Millipore clone Huts-4 #MAB2079Z), integrin β3 (#13166, CST), integrin β5 (#3629, CST), integrin ⍺6 (Millipore #MAB1378). Appropriate secondary antibodies (peroxidase-conjugated IgG, Bio-Rad) were used. The ECL Kits (ThermoScientific) were used for signal detection. The Westerns were imaged by Amersham Imager 600. Images were later imported to ImageLab software (Biorad).

### Drug treatments and IC50 calculations

For dose dependent drug treatment experiments, cells were plated onto 96 well plates (7.5×10^3^ per well) and were allowed to adhere. The next day media was changed to non-conditioned or CAF-CM with addition of 2% FBS in order not to disrupt cycling of the cells. For experiments treated with SGK inhibitor BLU6340, cells were pre-treated with BLU6340 (250 nM) prior to the administration of chemotherapy and spiked every 48h. The drugs were administered using the Tecan D300e digital dispenser with a range of drug concentrations corresponding to a log 1/3.16 step down from the highest indicated concentration. 72h after, end-point cell viability assays were performed with Presto Blue HS (Invitrogen), values were normalized to corresponding DMSO-treated control wells and plotted in GraphPad Prism software Boston, Massachusetts USA, www.graphpad.com. Cell response to drug concentrations was analysed using non-linear regression to fit the data to log(drug) vs. response (variable slope) curve. Obtained IC50 values for each treatment condition were presented on the graph.

### DNA damage foci Immunostaining, microscopy and quantification

Cells were plated on cover slips and once adhered, treated with the indicated chemotherapies either in the presence of 10% FBS, NC or CAF-CM for the duration of the experiment. Later, the cells were fixed for 20 min in ice-cold 90% methanol, rinsed in PBS, kept in blocking buffer (1×PBS, 5% goat serum, 0.3% Triton X100) for 60 min and further incubated with mouse yH2AX primary antibody (1:100, Millipore) overnight. The next day coverslips were incubated with corresponding secondary Alexa Fluor-conjugated antibodies (1:250, Thermo Fisher Scientific) at RT for 60 min. Coverslips were washed 3 times in PBS, DAPI was added in one of the PBS washes, and mounted using Shandon Immumount mounting media (9990402). The signal was visualized using either Zeiss LSM 880 Upright Confocal System, or Keyence BZ-X800 immunofluorescent microscopes. Image analysis, intensity measurements and foci counts were performed either manually, by FIJI/Image J^87^, or using modified speckle counting pipeline from Cell Profiler 4.0^88^.

### Comet assay

HPAC, SW1990, and SUIT-2 cells were plated on 6 wells (200 x10^3^/well). The next day media was changed to CAF-CM, and cells were treated with either gemcitabine 2 µM, SN-38 0.5 µM, or 5-FU 100 µM overnight. When indicated, cells were pre-treated with SGK1 inhibitor BLU6340 (250 nM for SW1990 and 500 nM for HPAC) for 24h, prior to the addition of CAF-CM with chemotherapies +/- BLU6340. Later, alkaline single cell electrophoresis was performed using Comet Assay High Throughput kit (Trevigen, R&D 4252-040-K) following manufacturer’s instruction. Briefly, cells were trypsinized, pellets were collected and washed twice in cold PBS. Cells were diluted in PBS w/o Ca^2+^ and Mg^2+^ and diluted 1:10 with low melting agarose. Drop of agarose/cell mix was spread on a comet assay slide, solidified at 4°C in the dark and submerged into lysis buffer overnight at 4°C. Next day comet slide was incubated in alkaline unwinding solution for 40 min at RT and subjected to electrophoresis at 17V for 34 min. After the run, cells were washed in distilled water and 70% ethanol, dried at 37°C for 6 hours and stained with SyBR Gold nucleic acid stain. Images were taken on Nikon Eclipse TE2000S fluorescent microscope using a 10× objective. Comets were evaluated using CometScore 2.0 free software and Olive Tail Moment^22^ of each individual cell was plotted using Graph Pad Prism Software, magenta lines portraying the median OTM value for each treatment. Mann-Whitney test was used for the evaluation of statistical significance.

### Matrix feeding assays, and Src and FAK inhibitor cell-based assays

Similarly, for yH2AX analysis, cells were plated on cover slips on Day 0, serum starved on Day 1 overnight, and the next day ECM proteins were added on the cells for 3h (FN1 20 µg/ml, collagen 1: 30 µg/ml, laminin 20 µg/ml, Matrigel 30 µg/ml, or RGD peptide 100 µg/ml), and the chemotherapy was added on and the cells were incubated for 24h (SN-38 250 nM, gemcitabine 0.5 µM). Cells were fixed with ice-cold methanol and stained for yH2AX foci (see above). RGD peptide was obtained from Selleck chemicals (#S8008).

Cells were starved O/N, and inhibitors were added 45 min before CAF-CM was added to the cells (Saracatinib 2 µM, Defactinib 2 µM). Three hours later cells were harvested. Matrix proteins were added at indicated concentrations and incubated on cells for 3h before harvesting lysates (Matrigel 3% vol/vol, Fibronectin [0.01 mg/ml], Collagen 1 [0.04 mg/ml], Laminin [0.01 mg/ml]). Matrix proteins were purchased from Corning (Matrigel, #354230) and Sigma (Laminin L2020, Fibronectin F1141, Collagen 1 C3867).

### Polar metabolite profiling by mass spectrometry

CAFs were serum starved for 4d as described above, the media collected spun at 1000×g for 10min, and metabolites extracted as described below. Quadruplicate samples were used for each condition. Non-conditioned DMEM media was used as a control. 10 µl media was collected and metabolites extracted with 90 µl extraction buffer (80% methanol, 25 mM ammonium Acetate and 2.5 mM Na-ascorbate prepared in LC-MS water, supplemented with isotopically labelled amino acid standards [Cambridge Isotope Laboratories, MSK-A2-1.2], aminopterin, and reduced glutathione standard [Cambridge Isotope Laboratories, CNLM-6245-10]). Samples were vortexed for 10 sec, then centrifuged for 10 minutes at 18,000g to pellet cell debris. Samples were directly injected (2 µl) into a ZIC-pHILIC 150 x 2.1 mm (5 µm particle size) column (EMD Millipore) operated on a Vanquish™ Flex UHPLC Systems (Thermo Fisher Scientific, San Jose, CA, USA). Chromatographic separation was achieved using the following conditions: buffer A was acetonitrile; buffer B was 20 mM ammonium carbonate, 0.1% ammonium hydroxide in water; resulting pH is around 9 without pH adjustment. Gradient conditions we used were: 0–20 min: linear gradient from 20% to 80% B; 20–20.5 min: from 80% to 20% B; 20.5–28 min: hold at 20% B at 150 uL/min flow rate. The column oven and autosampler tray were held at 25 °C and 4 °C, respectively. MS data acquisition was performed using a QExactive benchtop orbitrap mass spectrometer equipped with an Ion Max source and a HESI II probe (Thermo Fisher Scientific, San Jose, CA, USA) and was performed in positive and negative ionization mode in a range of m/z = 70–1000, with the resolution set at 70,000, the AGC target at 1 × 106, and the maximum injection time (Max IT) at 20 msec. HESI settings were: Sheath gas flow rate: 35. Aux gas flow rate: 8. Sweep gas flow rate: 1. Spray voltage 3.5 (pos); 2.8 (neg). Capillary temperature: 320°C. S-lens RF level: 50. Aux gas heater temp: 350°C.

### Data Analysis for metabolomics with TraceFinder

Targeted relative quantification of polar metabolites was performed with TraceFinder 5.1 (Thermo Fisher Scientific, Waltham, MA, USA) as previously described^89^ (referencing an in-house library of chemical standards (230 compounds and 35 isotopically labelled internal controls). Pooled samples and fractional dilutions were prepared as quality controls and injected at the beginning and end of each run. In addition, pooled samples were interspersed throughout the run to control for technical drift in signal quality. Data from TraceFinder was further consolidated and normalized with an in-house R script: (https://github.com/FrozenGas/KanarekLabTraceFinderRScripts/blob/main/MS_data_sc ript_v2.4_20221018.R) which performed technical and biological normalization and quality control (QC) steps. QC reported R-Squared (RSQ) (computed from pool dilutions) and the coefficient of variability (CV) (computed from pool reinjections). A cut off RSQ>0.95 and CV < 30% was set up for this study. Biological normalization was achieved by mean centering each metabolite that passed these QC thresholds and then using the average of these mean-centered values across each sample as a normalization factor. Normalized data was plotted in Excel and are included as Extended Data Table 1.

### Cell cycle analysis

SW1990 cells were plated on 6 cm plates and incubated with either non-conditioned or conditioned media from three patient derived CAF lines supplemented with 2% FBS for 24h. Cells were pulsed with 10 µM EdU for 1 hr before harvesting via trypsin digestion and fixation with 4% PFA and later subjected to Click chemistry using Click-iT™ EdU Pacific Blue™ Flow Cytometry Assay Kit according to manufacturer’s instructions (Thermo Scientific C10418). The percentage of cells in S phase was determined by the percentage of cells that incorporated EdU.

For the cell cycle synchronization studies SW1990 and AsPC cells were plated on 6 cm plates, treated with BLU6340 at 250nM when indicated, and subjected to a double thymidine block to synchronize cells at the G1/S border. Briefly, cells were incubated with 10% FBS DMEM containing 2mM thymidine for 18h, washed and incubated with media lacking thymidine for another 9h. Next, thymidine was added for another 18h. Cells later were released into thymidine-free media, time points were collected as indicated in figure legends. Similarly, cells were subjected to click chemistry (see above) followed by staining with PE anti-Histone H3 Phospho (Ser10) antibody (BioLegend 650808) to define the mitotic cell population. Later, cells were incubated with propidium iodide (Roche) and with 192 µg/ml RNase A (Sigma-Aldrich) for 30 min at RT before analysis to measure total DNA content. Data acquisition was performed using Beckman Coulter’s CytoFLEX LX Flow Cytometer at BIDMC Flow Cytometry Core and later analysed using FlowJo software.

### RPPA samples

BxPC-3, HPAC and SW1990 cells were plated in triplicates, serum starved overnight before CAF-conditioned media or DMEM media was added on the cells in the presence of 2 µM gemcitabine. Conditioned and control media was replaced every 24h, and at the day of harvesting (72h later), fresh conditioned media was added 1h before harvesting the lysates. Samples were lysed in RPPA lysis buffer, processed and analysed at MD Anderson core facility for 304 proteins and phospho-proteins as described previously ^29^. (And in https://www.mdanderson.org/research/research-resources/core-facilities/functional-proteomics-rppa-core.html). The raw values for the RPPA are included as Extended Data Table 2.

### Integrin blocking assays

SW1990 and HPAC cells were plated on glass bottom chamber slides, allowed to adhere, and starved overnight. Integrin beta 1 function blocking antibody (ITGB1 6S6 #MAB2253Z, 4 µg/100 µl), Integrin beta 4 function blocking antibody (Millipore #MAB2059 4 µg/100 µl) or control Mouse IgG1 antibody was added to the cells 3h before CAF-CM was added, and cells were incubated with CAF-CM for additional 3h before fixation with 4% PFA and immunostained for p-NDRG1 T346. Cells were imaged using identical acquisition settings for all the conditions. The images were analysed for fluorescence intensity in Fiji software^87^ using identical settings for all images. Results were plotted in GraphPad Prism.

### Plasmids, cloning, CRISPR-Cas9 KO, site-directed mutagenesis, and siRNA

LentiCRISPR v2 was a gift from Feng Zhang (Addgene plasmid # 52961)^90^, the constructs for NDRG1 KO were created by annealing the following sense and antisense oligonucleotides: KO-1s CACCGGTCTGCCACGTGGACGCCCC, KO-1as AAACGGGGCGTCCACGTGGCAGAC, KO-2s CACCGGTACCCCTCCATGGATCAGC and KO-2as AAACGCTGATCCATGGAGGGGTAC, KO-3s CACCG AGCAAATCGAGTTAGGATGT and KO-3as AAACACATCCTAACTCGATTTGCT, followed by subcloning in the LentiCRISPR v2 backbone. Constructs to create control cells were created by annealing the following sense and antisense oligonucleotides: one missense sequence targeting bacterial LacZ-s CACCTGCGAATACGCCCACGCGAT and LacZ-as AAACATCGCGTGGGCGTATTCGCA, and another targeting AAVS1 gene AAVS1-s CACCGGGGCCACTAGGGACAGGAC, and AAVS1-as AAACGTCCTGTCCCTCGTGGCCCC, were chosen as corresponding negative controls. Coding sequence of NDRG1 with mutated PAMs was ordered from GenScript, and further subcloned into pLenti6/V5-p53_wt p53 (gift from Bernard Futscher, Addgene plasmid # 22945)^91^ by cutting out the coding sequence of p53 and pasting the coding sequences of NDRG1 variants into via *XbaI* and *SpeI* restriction sites, allowing the generation of the C-terminal V5-tagged NDRG1 construct. The H194A mutation was introduced via site directed mutagenesis (QuikChange Lightning Multi Site-Directed Mutagenesis Kit, Agilent Technologies) using the following sense and antisense oligonucleotides H194A-s CCGGACATGGTGGTGTCCGCCCTTTTTGGGAAGGAAGAAATG, H194A-as oligo CATTTCTTCCTTCCCAAAAAGGGCGGACACCACCATGTCCGG. pLNCX chick-SRC was a gift from Joan Brugge (Addgene #13665), pBabe FAK-puro was a gift from Filippo Giancotti (Addgene #21155), empty plasmids were used as control (pLNCX and pBABE puro). pEGFP-N2-2XNLS-RNaseH1 delta 1-27 (WT) (Addgene #196702) and pEGFP-N2-2XNLS-RNaseH1 delta 1-27 (D210N) (Addgene #196703) were a gift from Karlene Cimprich^61^.

Transient knockdown of the NDRG1, SGK1, SGK2, SGK3, and SETX were achieved by transfections of cells with the smartpool of ON-TARGETplus Human NDRG1 siRNA (L-010563-00-0005), ON-TARGETplus Human SGK1 siRNA (L-003027-00-0005), ON-TARGETplus Human SGK2 siRNA (L-004673-00-0005), ON-TARGETplus Human SGK3 siRNA (L-004162-00-0005), ON-TARGETplus Human SETX siRNA (L-021420-00-0005), ON-TARGETplus Mouse Ndrg1 siRNA (L-045536-01-0005).

### Production of Lentiviruses and stable cell lines

For the generation of lentiviruses, 70-80% confluent HEK-293T cells were co-transfected with psPAX2 (a gift from Didier Trono, Addgene plasmid # 12260), pVSVg (gift from Joan Brugge), and one of the following vectors: LentiCRISPR v2 with subcloned g*NDRG1* KO-1,-2,-3, g*LacZ*, g*AAVS1* and pLenti6 wt *NDRG1*-V5 or pLenti6 H194A *NDRG1*-V5 by Lipofectamine 3000 (Thermo Fisher). Media was collected over a period of 24h to 72h post-transfection in 24h batches, pooled, centrifuged and filtered through a 0.45 µm filter. LentiCRISPR v2 infected cells were cultured in the presence of Puromycin (1 mg/ml) for 96h and the clonal cell populations were generated by fluorescence activated cell sorting (FACS). Later, clonal populations were checked for the successful knockout of NDRG1 and mixed to create an *NDRG1* KO mix. Clonal populations of cells generated from g*LacZ* and g*AAVS1* were mixed to create CNTR cells mixed population. *NDRG1* KO mix cells were later infected with either pLenti6 wt *NDRG1*-V5 or pLenti6 H194A *NDRG1*-V5 expressing viral particles, followed by the selection with Blasticidin (15 mg/ml) for 7-10 days.

For the generation of the stable doxycycline-inducible NDRG1 knockdown SW1990 cell lines, cells were transduced with viral particles (10^7^ TU/ml) containing SMARTvector Inducible human NDRG1 shRNA (V3SH7669-230407948, Horizon Discovery) and a scrambled control shRNA (VSC6570, Horizon Discovery) followed by puromycin selection at 1 µg/ml.

### Live Cell Imaging Assays

For real-time quantitative live-cell proliferation analysis, cells were seeded onto 96-well plates (5×10^3^ cells per well) in full media. The next day the plate was moved into the IncuCyte® S3 System (Essen BioScience) for live phase contrast recording of cell confluence. The confluence analysis was performed using the basic IncuCyte software settings and either raw confluency percentage or normalized confluency were used to present the data.

### Patient data

The Cancer Genome Atlas (TCGA), Pancreatic cancer cohort patient data (referred as PAAD in TCGA, but in this manuscript as PDAC) were obtained from cBioPortal^48,92,93^. Cloud-based application, TRGAted was used to examine patient survival based on pNDRG1 signal from RPPA data available for PDAC (PanCancerAtlas) dataset in TCGA. The code and processed data are open sourced and available on github and contain a tutorial built into the application^94^. Patient data from the Australian Pancreatic Genome Initiative (APGI) as part of the International Cancer Genome Consortium (ICGC) has been previously described^16,95^. Patients with resected oligometastatic disease were excluded. All patients were meticulously followed up. Patients with molecular data were followed up as per APGI protocol in keeping with standards of a prospective clinical trial. This included regular medical case notes review, contact with primary care providers and patients. Patients alive at the time of follow-up point were censored. The last follow-up period for patients still alive was October 2020. Kaplan-Meier survival analysis and log-rank rest were used to analyse disease-specific survival. For tissue resection stainings, PDAC tissue resections of a 59 y.o. male patient treated with 6 cycles of gemcitabine/Abraxane (Patient 1) and of the 37 y.o. female patient treated with 6 cycles of FOLFIRINOX and Stereotactic body radiation therapy (SBRT) (Patient 2) were used for p-NDRG1 staining^96^. The protocol was approved by the Dana-Farber Cancer Institute Institutional Review Board and informed consent was obtained from all patients enrolled.

## Bioinformatics

### ICGC RNAseq analysis

ICGC cohort sequencing reads were mapped to transcripts corresponding to ensemble 70 annotations using RSEM^97^. RSEM data were normalized using TMM (implemented in Biocondutor package ‘edgeR’)^98^, converted to counts per million and log2 transformed as previously described^16^. Genes without at least 1 c.p.m. in 20% of the sample were excluded from further analysis.

### ICGC Transcriptomic profiling and NDRG1 Expression

Transcriptomic profiling of the ICGC cohort was performed as part of the ICGC landmark study of PC^16^. Individual tumours were classified as squamous, pancreatic progenitor, aberrantly differentiated endocrine exocrine (ADEX) or immunogenic as previously described^16^. Normalized expression values for NDRG1 were converted to Z scores and stratified by subtype. Expression boxplots were created using the R package ‘ggpubr’^99^. Patients were dichotomized into 2 groups, based on high and low NDRG1 mRNA expression. The high and low groups were divided above and below the median.

### Tissue microarray analysis

Pancreas cancer tissue microarray (TMAs) PA805b (80 cases of PDAC/80 cores) with normal tissue as control, including TNM, clinical stage and pathology grade were purchased from US Biomax, Inc. TMA sections were processed with IHC protocol for the phospho-NDRG1 T346 (CST # 5482S,1:400) and total NDRG1 (Abcam, ab37897 1:400) followed by H&E counterstaining. Sections were evaluated by a board-certified anatomic pathologist (Dr. Raul S. Gonzalez) via the following criteria: the proportion of immunopositive cells in the field of view was assigned the number from 0 to 4 (0 <10%; 1: ≥10% to <25%; 2: ≥25% to <50%; 3: ≥50% to <75%; and 4: ≥75%). The staining intensity was categorized by relative intensity as follows: 0: no positivity; 1: weak; 2: moderate; and 3: strong. The analysis was performed using Fisher’s exact test using SAS Session software. P-value lower than 0.05 was considered significant. The sections were imaged using Olympus BX-UCB whole slide scanner equipped with Olympus UPLSAPO 20× and VS-ASW v2.7 software, and representative images were shown using HMS OMERO Service^100–102^.

### Tissue staining and immunohistochemistry

Resected tissues of the pancreatic cancer patients treated with a standard-of-care regimens (FOLFIRINOX or gemcitabine/Abraxane) in our institution^96^ were embedded in paraffin blocks, cut, deparaffinized in xylene and rehydrated in a descending ethanol series. Deparaffinized sections were subjected to antigen unmasking in SignalStain® Citrate Unmasking Solution (CST #14746) followed by blocking of nonspecific binding in TBST/5% normal goat serum solution. Later, sections were incubated with the primary antibodies against phospho-NDRG1 (#5482S, CST) and total NDRG1 (#9485S, CST), diluted 1:400 in SignalStain® Antibody Diluent (CST #8112) overnight at 4°C in a humidified staining tray. The next day, after the incubation with the secondary peroxidase-conjugated antibody, chromogenic detection was performed using the SignalStain® DAB Substrate Kit (#8059) for phospho-NDRG1 and for total NDRG1. For the hematoxylin/eosin (HE) staining the samples were stained for 2 seconds in Mayers’s Hematoxylin staining reagent (DAKO), rinsed in H2O, dehydrated in an ascending ethanol row and mounted with Permount Mounting Medium (Thermo Fisher, SP15500). Masson’s Trichrome staining was performed by BIDMC Histology Core. The sections were imaged using Olympus BX-UCB whole slide scanner equipped with Olympus UPLSAPO 20× and VS-ASW v2.7 software, and representative images were shown using HMS OMERO Service^100–102^. Patient resections were evaluated by a board-certified anatomic pathologist (Dr. Monika Vyas).

### Xenograft studies

All animal studies were performed according to protocols approved by the Institutional Animal Care and Use Committee at BIDMC. Athymic male nude mice (NU/J) were purchased from Jackson labs (#002019). SW1990 tumour cells (2×10^6^) and AsPC tumour cells (1×10^6^) were injected subcutaneously in one flank per mouse in 1:1 mix of Matrigel and PBS when the mice were 8-12 weeks old. In order to achieve NDRG1 knockdown for SW1990 tumour cells mice were fed Doxycycline chow (200 mg/kg) (Bio-Serv, S3888) prior to gemcitabine treatment once the tumours reached 200mm^3^. Tumour growth was monitored thrice weekly. Gemcitabine (25mg/kg) twice per week was administered to animals once the tumours reached 200mm^3^ in volume. Entire cohort of animals was sacrificed when tumour size of at least one animal was close to reach 1000mm^3^.

### Xenograft tissue staining

Xenograft tumour tissues were processed by BIDMC Histology core, paraffin embedded and cut, deparaffinized in xylene and rehydrated in a descending ethanol series. NDRG1 staining was performed as described above. For immunofluorescent staining deparaffinized sections were subjected to antigen unmasking in SignalStain® Citrate Unmasking Solution (CST #14746) followed by permeabilization in 1% Triton X100, and blocking of nonspecific binding in TBST/5% normal goat serum solution. Sections were later stained with mouse anti-Fibronectin (ab6328-250, Abcam), Pan Cytokeratin (AE1/AE3), Alexa Fluor™ 488 conjugated (53-9003-82, eBioscience) and rabbit phospho-NDRG1 (#5482S, CST) or Ki-67 (D3B5) Rabbit mAb (Alexa Fluor® 647 Conjugate) (#12075, CST). Appropriate secondary antibodies were used and whole section images were acquired using Olympus BX-UCB whole slide scanner equipped with Olympus UPLSAPO 20× and VS-ASW v2.7 software. Image analysis was performed in Qupath^103^.

### Spatial analysis of the phospho-NDRG1 and Ki67 signal intensity in relation to a modelled stromal border

A computational framework combining StarDist^104^, QuPath^103^, and Python^105^ was developed for the analysis of the whole tumour xenograft images stained for DAPI, Pan Cytokeratin, Fibronectin, as well as for phospho-NDRG1 or Ki67. StarDist in QuPath was used for deep learning-based cell nuclei detection, using the pre-trained ’dsb2018_heavy_augment.pb’ model for DAPI-stained nuclei in 2D fluorescence microscopy images. Cell boundaries were inferred with QuPath’s built-in cell expansion algorithm (5 µm radius). Nuclei with an area falling below the 5th percentile (around 15 µm²) were excluded. Cytokeratin, pNDRG1, and Ki67 positive cells were identified via QuPath built-in Random Forest model (“Random Tree”) trained on a manually curated representative cell set. Stromal borders were modelled to separate areas with high and low fibronectin content. High-fibronectin areas, modelled as stromal regions, were annotated in each image using a threshold-based pixel classifier.

Initial, manually determined, absolute threshold values from a subset of images were converted to their percentile values in their corresponding cell distribution, and then applied to the rest of the images by maintaining the same percentile value. This way, threshold values were fixed algorithmically thus minimizing subjective/operator bias. Best fit was determined using weighted least-squares. The fibronectin channel was blurred with Gaussian smoothing (σ = 10) before thresholding. Finally, the 2D signed distance (in µm) of each cell to its nearest stromal edge was calculated, with negative values indicating cells inside stromal regions. The class of each cell, along with its signal intensity measurements for the four channels, were exported for further data analysis and visualization in Python. Cells were grouped into bins, and within each fixed-size bin, the distance from stromal border was calculated as the midpoint of the bin position, and the signal intensity as the mean with its standard error. Results are presented as a mean distance from stromal border on the x-axis and the mean signal intensity on the left y-axis with a fixed bin size of 10 µm. The cell count of each bin is presented on the right y-axis. All QuPath scripts developed for this study, as well as Python notebooks used for visualization, are available at https://github.com/HMS-IAC/stroma-spatial-analysis.

### Isolation of Proteins on nascent DNA

The iPOND assay was performed as previously described^51^ with slight modifications. 400×10^6^ HPAC cells were pulsed with 10 µM EdU for 15 min. For a thymidine chase condition, cells were treated with 10 µM thymidine for 30 min, and hydroxyurea (HU) 4mM was added after the EdU pulse to stall replication forks. Cells were subsequently fixed using 1% formaldehyde for 20 min at room temperature. The crosslinking reaction was quenched using 1.25 M glycine and cells were washed three times with PBS. Cells were incubated with 0.25% Triton X-100 in PBS for 30 min at RT and pelleted. Permeabilization was stopped with 0.5% BSA in PBS. Cells were then pelleted again and washed with PBS. After centrifugation, cells were resuspended with the click reaction cocktail using biotin-azide (B10184, Thermo Fisher) and incubated for 2h at RT on a rotator. After centrifugation, the click reaction was stopped by resuspending cells in PBS containing 0.5% BSA. Cells were then pelleted and washed with PBS twice. Cells were resuspended in lysis buffer and sonicated. Lysates were cleared, then incubated with streptavidin-agarose beads overnight at 4^◦^C in the dark. The beads were washed once with lysis buffer, once with 1M NaCl, then twice with lysis buffer. To elute proteins bound to nascent DNA, the 2×SDS Laemmli sample buffer was added to packed beads (1:1, v/ v). Samples were incubated at 95^◦^C for 30 min, followed by immunoblotting.

### Proximity ligation assay

The PNCA-NDRG1 interaction and PCNA-RNAPII were detected with the Duolink PLA Kit (DUO92102, Sigma-Aldrich) according to the manufacturer’s protocol. For the PCNA-NDRG1 PLA, cells were plated on coverslips, grown for 24h, then treated for 16h either with 2mM hydroxyurea or PBS prior to incubation with CSK buffer (10 mM PIPES, 100 mM NaCl, 300 mM sucrose, 3 mM MgCl2, 1 mM EGTA and 0.5% Triton X-100™) for 10 min at 4°C and fixation with 4% PFA for 20 min at RT. Later, coverslips were permeabilized with blocking buffer. The samples were incubated with the primary rabbit NDRG1 polyclonal antibody (1:100, Cell Signaling) and mouse PCNA antibody (1:100 sc-271835, CST) diluted in blocking solution overnight at +4C. For the PCNA-RNAP II PLA cells were plated on coverslips, grown for 24h, underwent double thymidine block and released into 10% FBS media, fixed with 4% PFA for 20 min at RT. The samples were incubated with the primary rabbit PCNA polyclonal antibody (1:250, Abcam ab18197) and mouse RNAPII antibody (1:100, Sigma 05-623 clone CTD4H8) diluted in blocking solution overnight at +4C. After the washes with buffer A and B, incubation with bisbenzimide (1:5000, Hoechst stain, Sigma-Aldrich), cells were mounted. Microscopy was performed using a Zeiss LSM 880 Upright Confocal System, 63× PlanApo oil immersion objective and appropriate filter sets for Hoechst 405, Alexa Fluor 546, and Zen2009 software or Keyence BZ-X800 immunofluorescent microscopes. Confocal images were recorded in a Z-stack, further processed via ‘maximum intensity projection’ tool provided by Zen 2009 software. TRC PLA foci were quantified using modified speckle analysis pipeline from CellProfiler^87^.

### DNA fiber spreading

DNA fibers were performed as described previously^56^. Briefly, cells were sequentially pulsed with two thymidine analogues 50 µM CIdU (Sigma, C6891) and 150 µM (Sigma, 17125) with 2×PBS washes in between. When fork stalling was required, cells were pulsed with CldU, washed with PBS, treated with 2mM HU for 1hour, washed with PBS and pulsed with IdU. BLU6340 treatment was performed at 250nM either overnight or at 500nM for 30 min as indicated in figure legends. Pre-treatment with transcription inhibitors alpha-amanitin (10mg/ml) and flavopiridol (10 µM) was performed for 30 min prior to first pulse. Cells were then trypsinized and resuspended in PBS, 2.5 µL were pipetted on the top of SuperFrost plus slides (#48311-703, VWR). After 4 min, 7.5 µL spreading buffer (0.5% SDS, 200 mM Tris-HCl pH 7.4, 0.5 mM EDTA) was mixed with the cells for additional 2 minutes. Two glass slides were made per condition per experiment. Slides were tilted at 15 degrees to allow DNA fibers to run down the glass slide. Later, the fibers were air dried and then fixed in 3:1 methanol:acetic acid solution for 2 min, followed by the 2.5M HCl treatment for 30 minutes and 3% BSA/PBST blocking for 1 hour. Primary antibody incubation was performed for 1h with anti-CIdU (ab6326 Abcam, 1:100) and anti-IdU (BD-347580, 1:20). Following three washes with PBS, fibers were stained with appropriate secondary Alexa-Fluor conjugated antibodies for 30 minutes, washed, air-dried and mounted. Slides were imaged with Zeiss LSM 880 Upright Confocal System, 40× or 63× PlanApo oil immersion objective. Measurement of replication structures was performed manually using Fiji^87^. At least 200 fiber tracks were quantified per experimental condition per assay.

### RNA-DNA hybrid immunofluorescence

Catalytically inactive D210N human RNase H1 (dRNH1)^61^ was expressed as a GFP fusion in E. Coli BL21(DE3), induced using 200 μM IPTG overnight at 16°C, purified by sequential Ni-NTA affinity, heparin affinity, and gel filtration, and the monodisperse fraction was concentrated to 7 mg/ml^106^. Cells were plated on coverslips, fixed with ice-cold methanol for 30min and incubated with GFP-dRNH1 in 3%BSA (1:500) for 3h at 37C followed by two washes in PBS. Anti-DNA-RNA Hybrid S9.6 antibody (1:100, Kerafast ENH002) was used overnight at 4°C. After PBS washes, DAPI was added in one of the PBS washes, and mounted using Shandon Immumount mounting media (9990402). The signal was visualized using either Zeiss LSM 880 Upright Confocal System, or Keyence BZ-X800 immunofluorescent microscopes. GFP nuclear intensity measurements were performed using CellProfiler 4.0.

### HR reporter system

*Escherichia coli* Tus/*Ter* Replication Fork Barrier system was previously described^107^ to enable the analysis of mammalian stalled fork metabolism and repair. Briefly, a chromosomally integrated array of six 23 bp *Ter* sites mediates Tus-dependent, locus-specific replication fork stalling and HR on a mammalian chromosome, enabling direct quantitation of the repair products of mammalian replication fork stalling. To detect stalled fork repair events, the affinity of Tus protein for the Ter DNA element is used. Tus binding to 6-Ter-elements on the chromosome imposes a physical barrier to replication fork progression and the Tus/Ter fork stalling induces HR repair. This system has revealed that the HR at stalled forks (induced by Ter expression vector) differs from the DNA repair taking place at conventional double-strand break (DSB) (induced by I-SceI endonuclease expression vector). HR repair products of Tus-Ter-induced fork stalling can be detected by the means of FACS. The reporter contains two non-functional alleles of GFP (grey boxes) flanking synthetic exons encoding RFP in a tail to head orientation. If DSBs or stalled forks are repaired by HR, the truncated GFP allele may undergo a gene conversion event, copying a short region of the 5’-truncated GFP allele off the intact sister-chromatid, producing a wild-type GFP allele (Green box, GFP+ cells). These short-tract gene conversion (STGC) events (GFP+) are the most frequent repair event. A distinct minority of HR repair termed long tract gene conversion events (LTGC), error-prone repair, convert the mutant to wild type GFP (GFP+) and duplicate the entire interior of the reporter cassette, which includes synthetic exons encoding RFP (Green box and Red oval). LTGC events create cells that are both GFP+/RFP+. Later, three repair events might occur: 1) short tract gene conversion of GFP (STGC), which appears to be the majority of HR events in mammalian cells, 2) long tract gene conversion (LTGC) of GFP accompanied by cassette duplication and RFP expression, error prone HR outcome or 3) tandem duplications (a mutagenic repair outcome specific to replication fork stalling which occurs without successful HR without gene conversion of mutant GFP). Depending on the ratio of GFP+RFP-/GFP+RFP+ cells, the impact of the NDRG1 knockdown on DSB repair was evaluated. Detailed schematics presented in Extended Data Figure 5.

### CellProfiler analysis

Publicly available CellProfiler^88^ ’Speckle Counting’ pipelines were modified according to nuclei/speckle size and used for yH2AX and TRC foci counts. Modified ‘Human C-N translocation’ pipeline was used for the measurement of the Nuclear, cytoplasmic, and whole cell intensities of the assayed markers.

### Statistical analysis

GraphPad PRISM 10, R software, Python, and Tableau were used for statistical and visual analyses. Sample size and error bars are reported in the figure legends. Exact P values are shown where possible. Unless otherwise noted, statistical tests were performed using unpaired two-tailed Student t-test. The number of times experiments were performed with similar results is indicated in each legend. P-values less than 0.05 were considered significant.

## Data Availability Statement

CAF lines and Addgene plasmids are under materials transfer agreements. Data in Figure 3 and Extended Figure 3 are from 32 combined TCGA PanCancer Atlas studies (https://www.cbioportal.org), Pancreatic Adenocarcinoma (TCGA, PanCancer Atlas) (https://www.cbioportal.org/study/summary?id=paad_tcga_pan_can_atlas_2018) and ICGC PDAC datasets, which have been published. ICGC data have been deposited in the European Genome-phenome Archive (EGA): accession code EGAS00001000154.

## Code Availability Statement

All QuPath scripts developed for this study, as well as Python notebooks used for visualization, are available at https://github.com/HMS-IAC/stroma-spatial-analysis.

Targeted relative quantification of polar metabolites was performed with TraceFinder 5.1 (Thermo Fisher Scientific, Waltham, MA, USA) and data from TraceFinder was further consolidated and normalized with an in-house R script available at GitHub which performed technical and biological normalization and quality control (QC) steps. https://github.com/FrozenGas/KanarekLabTraceFinderRScripts/blob/main/MS_data_scri pt_v2.4_20221018.R)

## Acknowledgements

We thank members of the Muranen lab and Cancer Research Institute (BIDMC, Boston, MA) for helpful discussions. Our warmest gratitude extends to Dr. Manav Gupta (Harvard PhD program in Biological & Biomedical Sciences (BBS), Stem Cell Program, Division of Hematology/Oncology and Division of Pulmonary Medicine, Boston Children’s Hospital, Boston, MA) for providing expertise with Comet assay and DNA fiber assay and to Caroline Fahey, M.Sc, (Harvard BBS program) for technical assistance with fiber stainings during pilot experiments. We also thank Samuel Lee and Joseph Abirached from Preclinical Murine core at BIDMC for help with mouse experiments, Sebastien Meurant (Department of Biology, University of Namur, Namur, Belgium) for assistance with experiments in the beginning of the project. We also warmly thank Dr. Costas Lyssiotis (Rogel Cancer Center, University of Michigan Medical School, Ann Arbor, MI) for helpful discussions, Dr. Sarah Robyn Wessel (Department of Biochemistry, Genome Maintenance Program in the Vanderbilt-Ingram Cancer Center, Vanderbilt University School of Medicine, Nashville, TN) for helpful advice with iPOND experimental setup; we thank Dr. Nicholas Borcherding (Department of Pathology and Immunology, Washington University School of Medicine, St Louis) for his help with TRGAted application, and Dr. Nada Kalaany (Boston Children’s Hospital, Boston, MA) for materials and helpful discussions. In addition, we thank Dr. Esin Işik (Dana Farber Cancer Institute, Boston, MA) for useful advice on dRNH1 staining and valuable suggestions. We thank Bryant D Miller (Boston Children’s Hospital, Boston, MA) for excellent technical assistance in purifying recombinant dRNH1. We apologize to all authors whose work could not be discussed due to space constraints.

This work was supported by Finnish Cultural Foundation (Suomen Kulttuurirahasto) Postdoc pool fellowship 2019 and 2020 to N.K., H.M.D. and K. A. C. by HMS graduate school (BBS), and K.A.C. by the NSF DGE 2140743, NCI 3R35CA242428 and Black in Cancer/Emerald Foundation Career Transition Award to M.U.J.O., Cancer Research UK clinical training ward and post-doctoral research bursary for S.D., The National Health & Medical Research Council of Australia, The Cancer Council NSW, Cancer Institute NSW, Royal Australasian College of Surgeons, The Avner Pancreatic Cancer Foundation, St Vincent’s Clinic Foundation and R. T. Hall Trust (S.D., R.UG, D.K.C), BCH BTREC to N.K. who is a Pew Scholar, Janne and Aatos Erkko Foundation to S.R. and P.K., NIH R37CA283575, V Scholar Award V2021-026, National Pancreas Foundation Research Grant NPF-5615796, Sky Foundation Research Grant to C.J.H., R01CA74305 to P.A.C., HMS Foundry Award Program to A.A.R and S.F.N., R35CA263813 and R01GM134425 to R.S., American Cancer Society grant #RSG-19-0201-CSM and NIH/NCI grants #R01CA258372 and #R00CA180221 to T.M.

## Author contributions

N.K. planned the study, designed and performed experiments, analysed the data, and wrote the manuscript

K.A.C. performed the experiments, helped with mouse work and analysed the data

H.M.D., performed the experiments and analysed the data

A.A.R. performed spatial whole image analysis and analysed the data

N.A.W. performed the Tus/Ter reporter experiments and analysed the data

R.G. analysed and scored the NDRG1 IHC data

M.V. analysed the NDRG1 IHC data

B.T., S.M.H performed experiments

L.M.S. performed the statistics analysis for the IHC data scoring

S.D., R.U.G., D.K.C. analysed the ICGC patient data

K.L.F., S.W., J.C. provided BLU6340 compound and expertise

M.U.J.O. Helped with live cell imaging and tissue culture experiments

M.S.T. provided dRNH1 and expertise on dRNH1 staining

V.M., S.R., P.K. provided expertise on NDRG1 structure and function

P.M. assisted with statistical analyses and analysed RPPA data

A.K.B., Z.J., and C.J.H. isolated and provided the CAF Patient 1, CAF Patient 2 lines and expertise in CAF-CM experiments

J.E.G. assisted with patient data discussion

R.H. provided the CAF line

J.G.C. helped design and execute the animal work

B.P. performed polar metabolite profiling by mass spectrometry and provided expertise

N.K. provided expertise on metabolomics

P.A.C. provided expertise and assisted with data discussion

S.F.N supervised spatial image and data analysis

R.S provided expertise with Tus/Ter reporter experiments, provided expertise and assisted with data discussion

T.M. supervised, planned and designed the study, analysed the data, and wrote the manuscript.

## Materials & Correspondence

All correspondence should be addressed to Dr. Taru Muranen

## Extended Data Figure legends

**Extended Data Fig.1.**
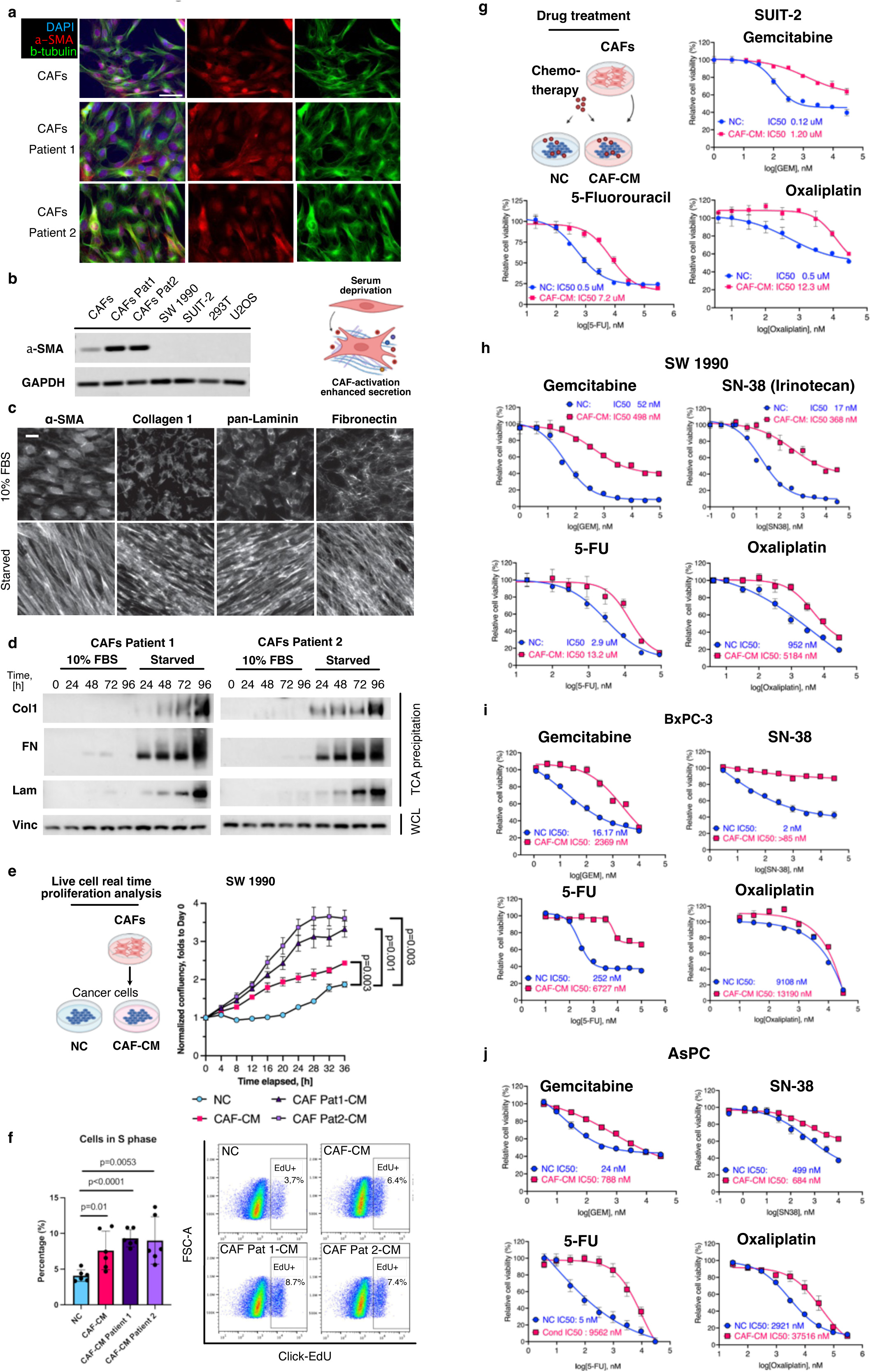
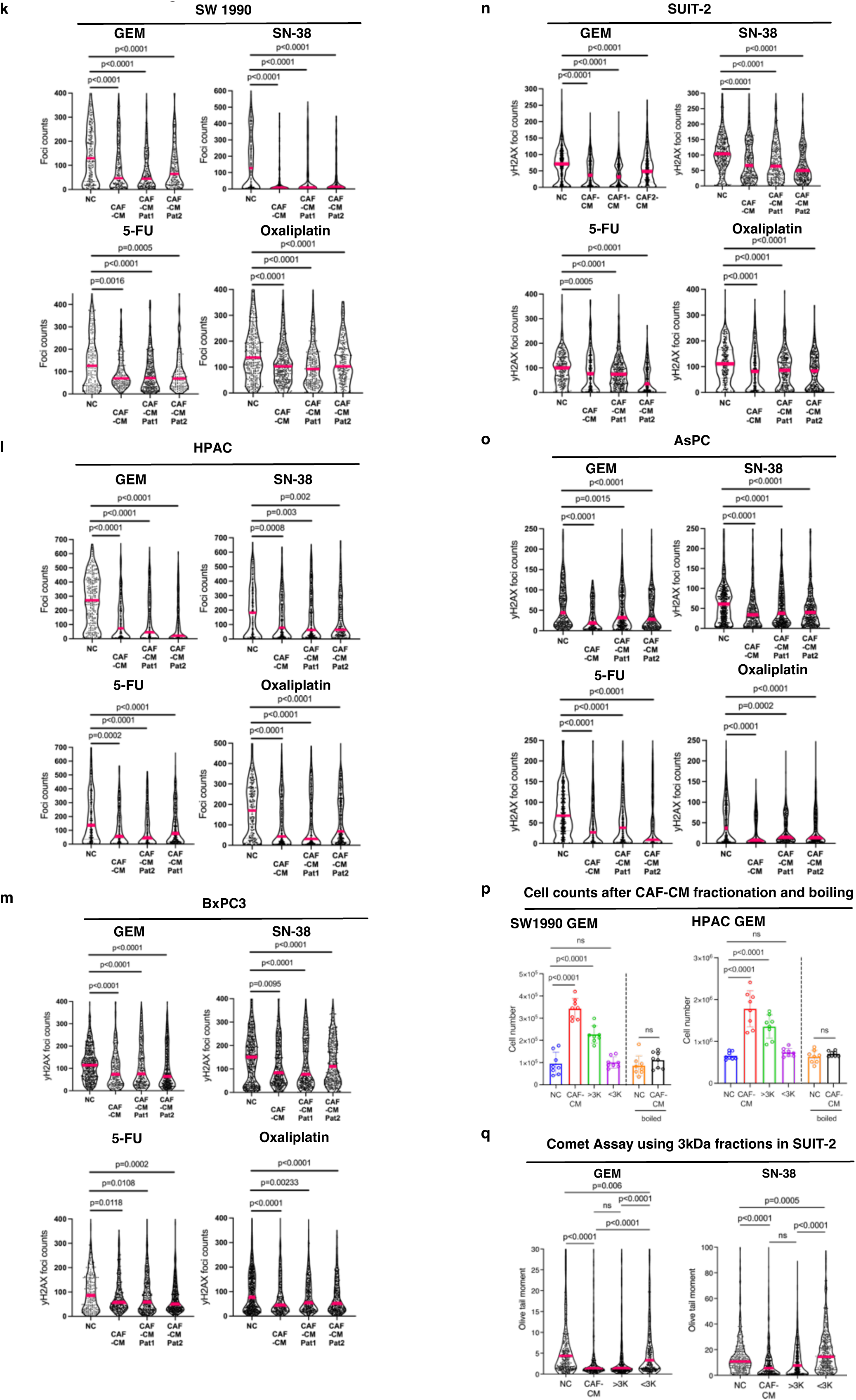
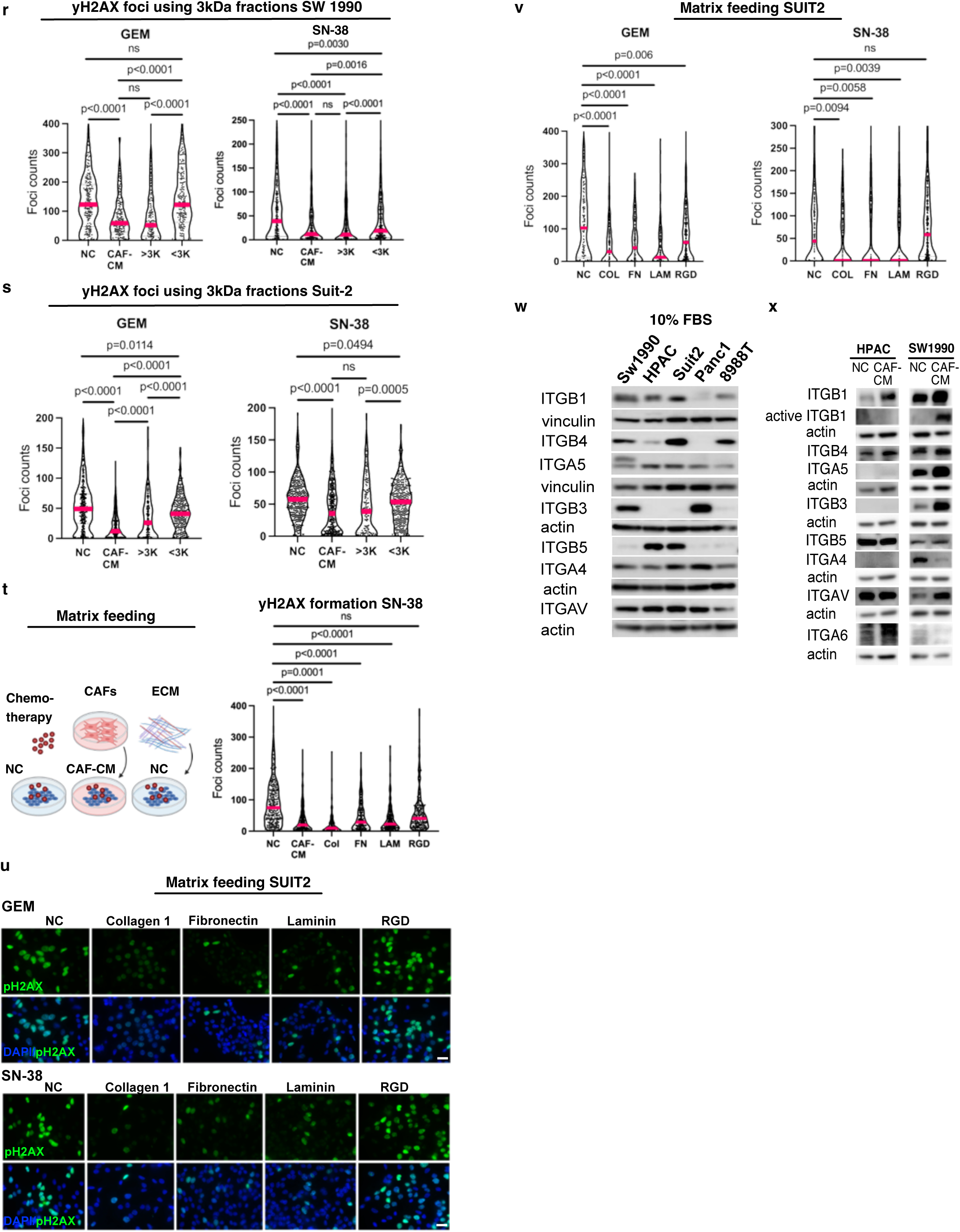
Starvation stimulates cancer-associated fibroblasts (CAFs) to increase secretion of ECM proteins, which promotes drug resistance in pancreatic cancer cells. **a**, To characterize the PDAC CAF lines used in this study, three lines were stained and imaged by IF for α-SMA, b-tubulin and DAPI. Scale bar: 50µm. **b**, The same CAF lines as above, and indicated epithelial cell lines were probed for α-SMA and GAPDH by Western blot. **c**, Deposition of Collagen 1, pan-Laminin, and Fibronectin by starved CAFs was visualized by immunofluorescence at Day 9, α-SMA was used as a marker for CAF activation, scale bar: 20µm. **d**, Additional CAF lines were kept in 10% serum, or starved similarly as in Fig. 1a, and media were harvested at indicated timepoints, TCA precipitated and probed for Collagen 1 (Col1), Fibronectin (FN) and pan-Laminin (Lam). The whole cell lysates (WCL) were probed for Vinculin. The TCA precipitates were loaded according to protein concentrations in the WCL. **e**, CAF-CM effect on SW1990 cell proliferation was analysed using automated live cell imager over 36h. Three CAF lines were used and NC media was used as a control. Data are represented as the confluency fold change normalized to day 0. Error bars represent SEM. Data were analysed by Mann-Whitney test. **f**, CAF-CM effect on SW1990 cell cycle was analysed using flow cytometry with cells labelled with Click-EdU (S-phase). CAF-CM increases PDAC cells in S-phase. Data are shown as mean with SD. Student’s t-test test was used for significance. **g**, SUIT-2 tumour cells’ IC50 responses towards chemotherapies were compared in CAF-CM (magenta) and in non-conditioned media (NC, blue). Fitted dose response curves to gemcitabine, 5-fluorouracil, and oxaliplatin were used to generate IC50 values for each treatment condition. Error bars are SEM corresponding to technical replicates. **h-j**, SW1990, BxPC-3 and AsPC tumour cell IC50 responses towards chemotherapies were compared as above (**g**) in CAF-CM (magenta) and in NC media (blue). Fitted dose response curves to gemcitabine, SN-38, 5-fluorouracil, and oxaliplatin were used to generate IC50 values for each treatment condition. **k**, SW1990 cells were treated with 1µM gemcitabine, 0.5 µM SN-38, 100 µM 5-FU, or 50 µM oxaliplatin for 24h in NC or CAF-CM (obtained from three patient-derived CAF lines). yH2AX foci analysis was used to assess DNA damage. Cells were fixed and immunostained for yH2AX followed by image acquisition and quantification by CellProfiler, data are shown as foci counts per single nucleus. >200 nuclei were analysed per condition. Mann-Whitney test was applied, magenta bars represent median. **l-o**, Similarly, as above in (**k**), HPAC, BxPC3, SUIT-2 and AsPC cells were analysed for yH2AX foci in three different CAF-CM vs. NC media, treated with 1µM gemcitabine, 0.5 µM SN-38, 100 µM 5-FU, or 50 µM oxaliplatin. **p**, CAF-CM and NC media were either boiled, or CAF-CM was filtered using 3kDa cut-off spin filters. PDAC cells (SW1990, HPAC) were treated with indicated medias (NC, CAF-CM, CAF-CM <3kDa, CAF-CM >3kDa, NC boiled, CAF-CM boiled) for 24h in the presence of 1 µM (SW1990) or 2 µM (HPAC) gemcitabine and cell numbers were counted. Student’s t-test was used to measure significance, data are shown as mean with SD. **q**, SUIT-2 were analysed for DNA damage by the alkaline comet assay. Proteins were filtered from CAF-CM using 3kDa cut-off spin filters. Cells were treated with 1µM gemcitabine or 0.5µM SN-38 for 24h in NC, CAF-CM, filtered CAF-CM (<3kDa proteins) and the top fraction (>3kDa proteins). Afterwards cells were plated on Comet assay slides and DNA was run under alkaline conditions and stained with SYBR-gold. Images were acquired and quantified using Comet Score software. Olive Tail Moment (OTM) were plotted in Graphpad Prism8 and Mann-Whitney test was used to establish significance. Magenta lines are median, >200 cells were analysed for each treatment condition. **r-s,** SW1990 and SUIT-2 cells were analysed for yH2AX foci in 1 µM gemcitabine or 0.5 µM SN-38-treatment after incubation for 24h in the different fractionated media: NC, CAF-CM, CAF-CM <3kDa, or CAF-CM >3kDa. yH2AX foci were measured and quantified as above (**k**). **t**, SW1990 cells were starved overnight after which NC, CAF-CM, purified matrix proteins (collagen 1, fibronectin, laminin) or RGD peptides were added to the cells along with 0.25 µM SN-38 for 24h. yH2AX foci were quantified as above (**k**). Mann-Whitney was used to measure statistical significance, and magenta lines represent median. **u**, SUIT-2 cells were treated as above with 0.5 µM gemcitabine or 0.25 µM SN-38 with indicated media or purified matrix proteins, stained for yH2AX (green) and DAPI (blue) and imaged. Representative images are shown. Scale bar: 20 µm. **v**, yH2AX foci analysis of SUIT-2 cells treated with NC, CAF-CM, or purified matrix proteins in the presence of 0.5 µM gemcitabine (GEM) or 0.25 µM SN-38 for 24h. yH2AX foci were quantified as above (**k**). Mann-Whitney was used to measure statistical significance, magenta lines represent median. **w,** Integrin profiling of PDAC lines. SW1990, HPAC, SUIT-2, PANC-1 and PA-TU-8988T cells were grown in 10% serum and probed for indicated integrins by Western blotting. Vinculin/actin was used as a loading control. **x**, CAF-CM changes integrin expression profile. HPAC and SW1990 cells were grown in NC or CAF-CM and probed for indicated integrins by Western blot. Actin was used as a loading control.

**Extended Data Fig. 2.**
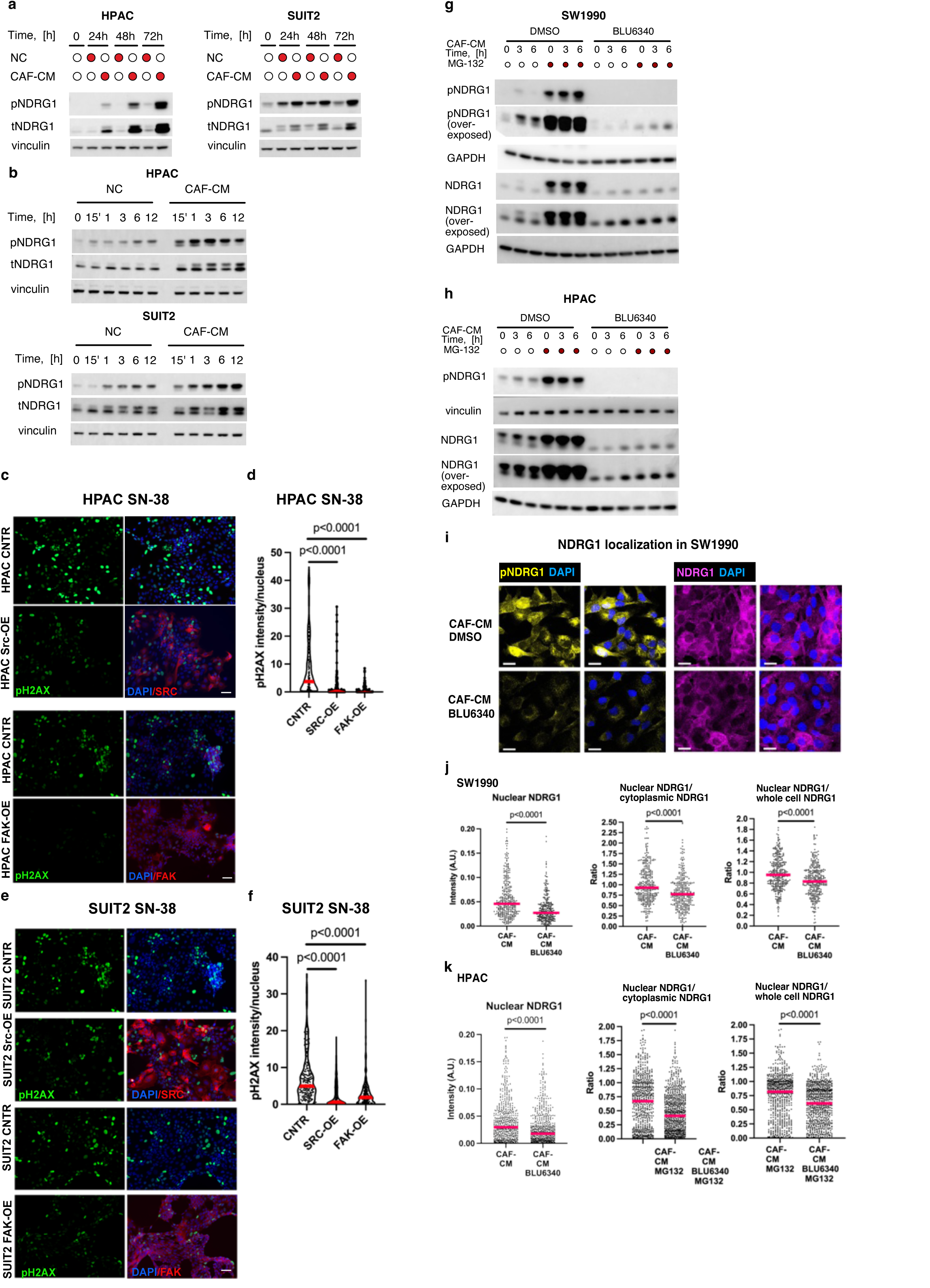
Inhibition of SGK1 abolishes NDRG1 phosphorylation in response to CAF-CM and sensitizes PDAC cells to chemotherapy. **a-b,** HPAC and SUIT-2 cell lines were treated with NC or CAF-CM for the indicated times (**a**: 24-72h, **b**: 0-12h) and cells were probed for p-NDRG1, t-NDRG1 and vinculin to assess phospho- and total NDRG1 levels over time. **c-f**, SRC or FAK kinases were overexpressed in PDAC cells (HPAC, SUIT-2) and cells were treated with 0.5 µM SN-38 for 24h. Cells were fixed and immunostained for yH2AX (green), SRC/FAK (red), and nuclei were counterstained with DAPI (blue). Scale bar: 50µM. yH2AX foci were quantified by CellProfiler, data are shown as yH2AX intensity/nucleus. >100 nuclei were analysed per condition. Mann-Whitney test was applied, magenta bars represent median. **g**,**h** SW1990 and HPAC cells were serum started overnight, CAF-CM was added for indicated times +/- BLU6340 (250nM), and +/- MG132 (10µM) to inhibit proteasome function. Cells were lysed and probed for phospho- and total-NDRG1, vinculin and GAPDH were used for loading. **i**, SW1990 cells were plated on coverslips, starved overnight, pre-treated with DMSO or SGK1 inhibitor (250nM BLU6340) for 30 min and treated with CAF-CM +/- BLU6340 for 6h. Cells were fixed and imaged for phospho-NDRG1 (yellow) and total NDRG1 (magenta), nuclei were counterstained with DAPI (blue). Scale bar: 20 µm **j,k**, Images from SW1990 and HPAC cells treated as described in (**i**) were analysed using CellProfiler to measure whole cell, nuclear, or cytoplasmic NDRG1. NDRG1 intensity was measured to evaluate whether phosphorylation affects NDRG1 localization. Nuclear intensities and ratios of nuclear intensities to either whole cell intensities or to cytoplasmic intensities were plotted in GraphPad Prism10, magenta lines are medians, Mann-Whitney test was used to establish significance. For better data visualization of NDRG1 localization in HPAC, cells were treated with MG132 to prevent protein degradation for the duration of the treatment. At least 330 cells were analysed per condition.

**Extended Data Fig. 3.**
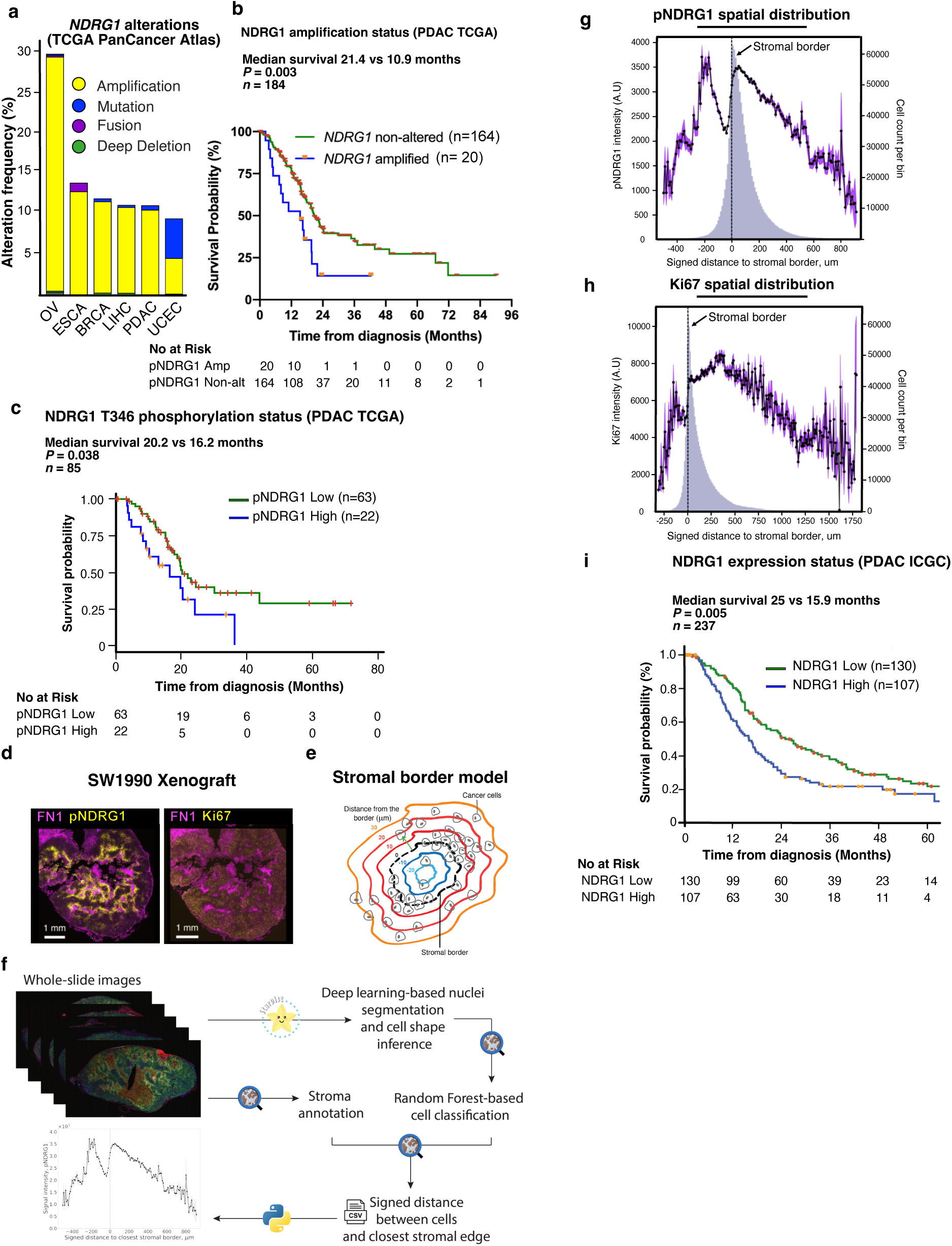
NDRG1 is highly expressed in pancreatic cancer and expression correlates with poor survival. **a,** NDRG1 data from The Cancer Genome Atlas (TCGA) project from cBioPortal. Seven cancer types where *NDRG1* is most frequently altered are shown. *NDRG1* is amplified in ∼10% of PDAC (OV: ovarian, ESCA: Oesophageal, BRCA: breast, LIHC: liver, PDAC: pancreas, UCEC: uterine corpus endometrial carcinoma). **b**, Survival probability of human PDAC patients (TCGA) with either amplified or non-amplified *NDRG1* is shown. Log-rank test was used for significance. **c**, Survival probability of human PDAC patients (TCGA) project based on RPPA analysis of the NDRG1 phosphorylation status. Log-rank test was used. **d-h**, Details for the spatial image analysis. **d**, Whole tumour image of SW1990 gemcitabine-treated xenograft stained for Fibronectin (magenta) and pNDRG1 (in yellow) and for Fibronectin (magenta) and Ki67 (in yellow) used in Fig. 3e. Scale bars are 1mm. **e**. Stromal border model used for spatial image analysis. The 2D signed distance (in µm) of each cell to its nearest stromal edge was calculated, with negative values indicating cells inside stromal regions. The black dotted line indicates the stromal border. Cells positive for a marker of choice were grouped into bins depending on their distance from closest stromal border. **f**, Image analysis workflow to study the spatial organization of pNDRG1 signal intensity in relation to the stromal border, combining deep learning for cell detection using a pre-trained StarDist model, training a QuPath Random Forest model for cell classification, threshold-based pixel classifiers for stroma annotation, 2D signed distance as spatial information, and Python for data analysis and visualization. **g**,**h,** Spatial analysis of pNDRG1 and Ki67 signal intensity in cancer cells given their signed distance to the closest stromal border (Distance in µm is on x-axis). Negative distance indicates cells located within stromal regions, while positive distances show cells located outside of these regions. Cells positive for pNDRG1 or Ki67 were grouped into bins (in violet) based on distance from signed stroma border and cell number per bin is plotted on the right y-axis, bin size is 10 µm. Purple sensogram represents an average maximum cellular signal intensity of pNDRG1 or Ki67-positive cells (black dots; Standard Error of the Mean (SEM) indicated as purple overlays) per bin and values are plotted on the left y-axis. pNDRG1 positive cells were distributed along -500 µm to 1000 µm in relation to closest stromal border; Ki67-positive cells were distributed along -300 µm to 2000 µm in relation to closest stromal border. Note highest positive pNDRG1 or Ki67 cell number per bin occurs in close proximity to the stromal border, while the amount of pNDRG1 and Ki67-positive cells declines the further those cells are away from one. **i,** Survival probability of human PDAC patients with either high or low NDRG1 mRNA expression. NDRG1 mRNA expression status in the ICGC PDAC cohort was analysed (high vs low) and overall survival was plotted over time. Kaplan-Meier survival analysis and log-rank rest were used to analyze disease-specific survival.

**Extended Data Fig. 4.**
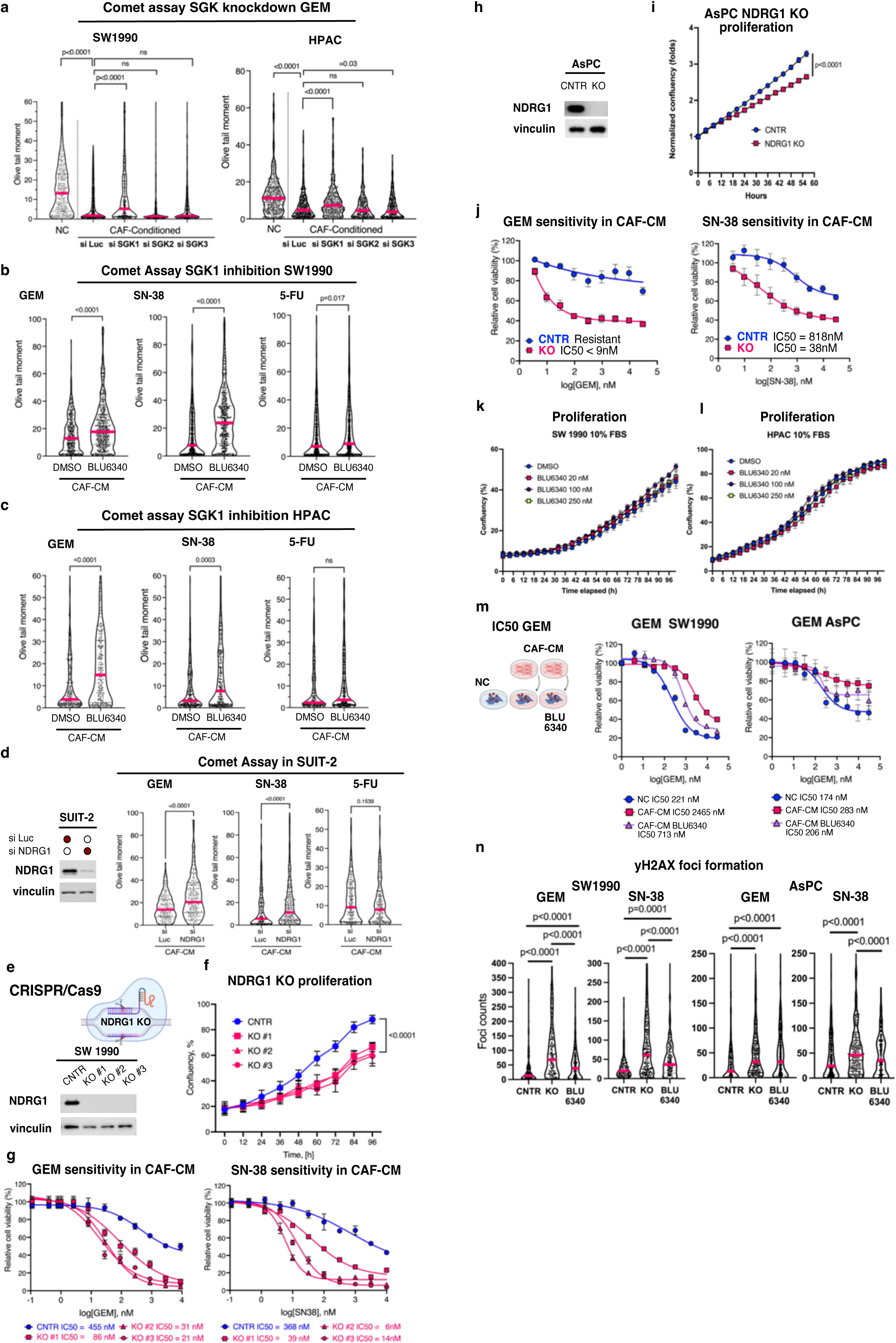
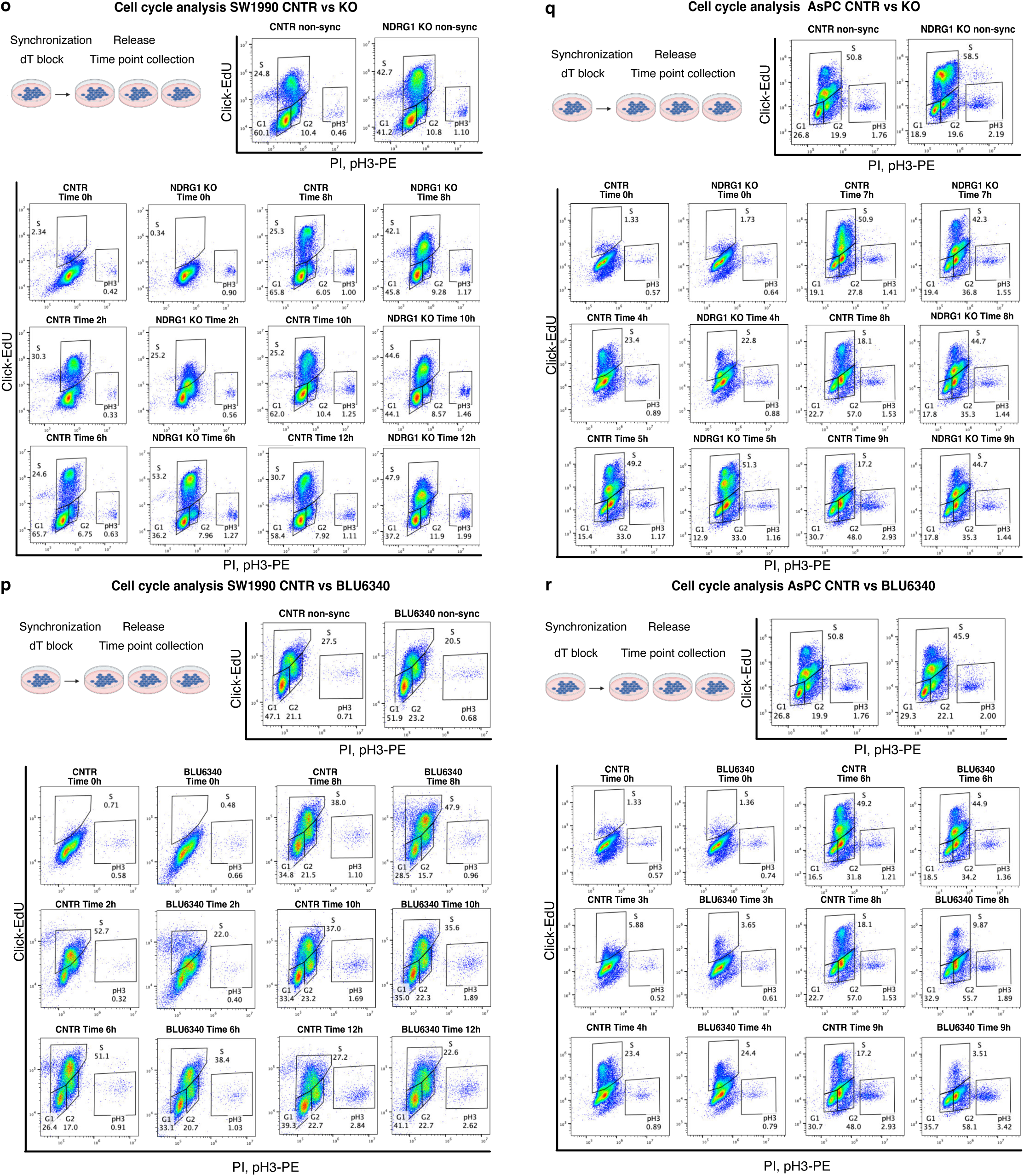
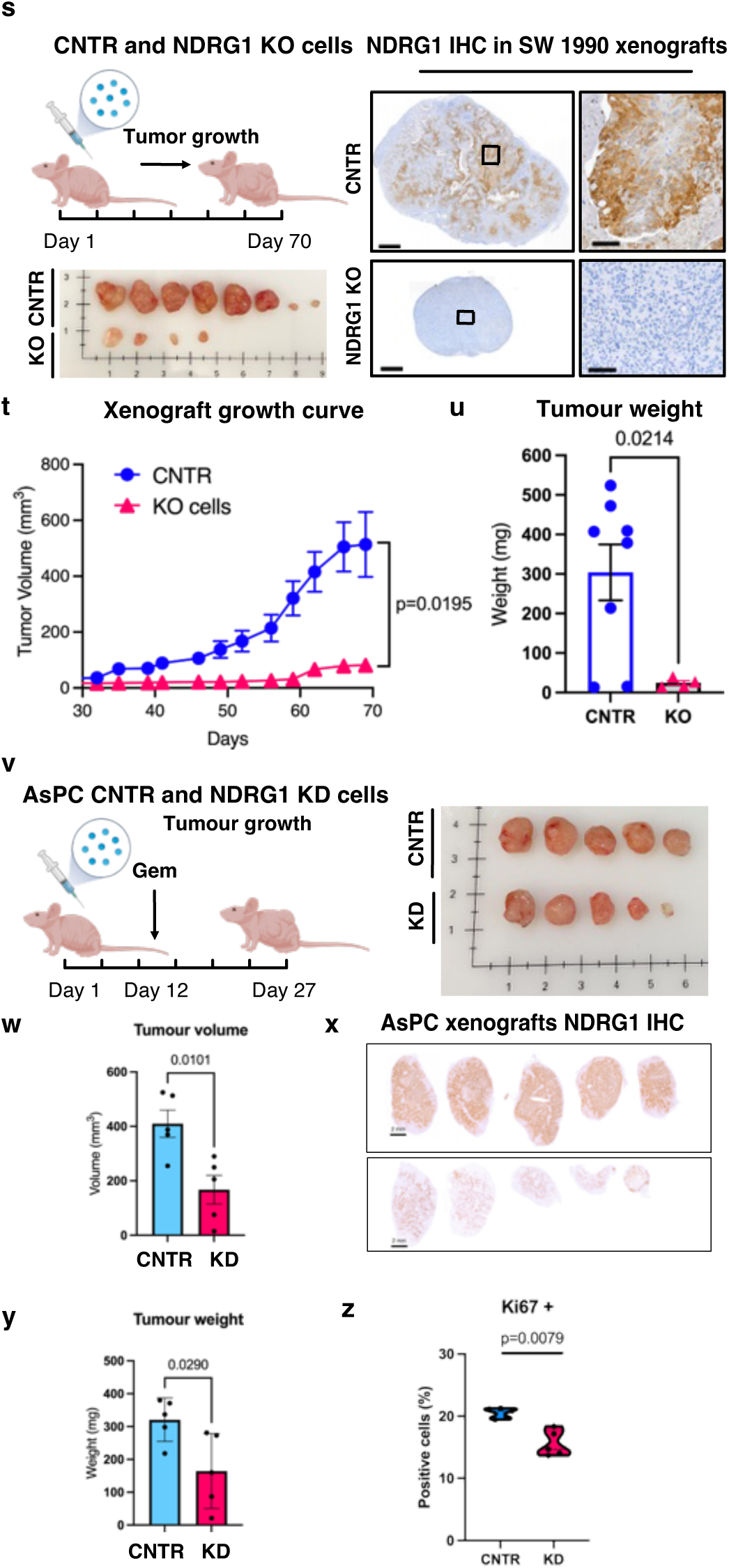
Loss of NDRG1 leads to higher levels of DNA damage in response to chemotherapies. **a,** SW1990 and HPAC cells were transfected with short interfering RNA (siRNA) against SGK1, SGK2, or SGK3 and treated with 1µM gemcitabine in NC or CAF-CM and analysed by Comet assay, OTM values were plotted, and two-tailed Mann-Whitney test was used for analysis. Magenta lines represent median values. >250 cells were analysed. **b**,**c** SW1990 and HPAC cells were treated with 1 µM gemcitabine, 0.5 µM SN-38, and 100 µM 5-FU in CAF-CM +/- 250 nM SGK1 inhibitor BLU6340, 500nM for HPAC and 24h later subjected to Comet assay. OTM values were analysed using two-tailed Mann-Whitney test. Magenta lines represent median values. >250 cells were analysed. **d,** SUIT-2 cells were transfected with short interfering RNA (siRNA) against NDRG1 and subjected to immunoblotting to verify the knockdown. The cells were treated with 1 µM gemcitabine, 0.5 µM SN-38, or 100 µM 5-FU in CAF-CM and analysed by Comet assay. Comet assay images were quantified, OTM values were plotted, and two-tailed Mann-Whitney test was used for analysis. >175 cells were analysed, magenta lines represent median values. **e**, SW1990 *NDRG1* KO cells were created using CRISPR/Cas9 system. Data from 3 clonal cells lines are shown, cells used as control (CNTR) are described in Material and methods. **f**, SW1990 CNTR cells (blue) and *NDRG1* KO (magenta) cells were subjected to live image proliferation analysis. Growth curves of cells in 10% FBS are shown over 96h, ANOVA was applied for analysis, error bars represent SEM. **g,** Fitted dose response curves for gemcitabine and SN-38 in CAF-CM were used to generate IC50 values for SW1990 *NDRG1* KO (magenta) and CNTR (blue) cells. Error bars are SEM corresponding to technical replicates. **h**, *NDRG1* was knocked out in AsPC cells using CRISPR/Cas9 and knock-out was verified by Western blotting for NDRG1. **i**, AsPC CNTR cells (blue) and *NDRG1* KO cells (magenta) were subjected to live image proliferation analysis. Growth curves of cells in 10% FBS are shown over 96h, Student’s t-test was applied for analysis, error bars represent SEM. **j,** Fitted dose response curves for gemcitabine and SN-38 in CAF-CM were used to generate IC50 values for AsPC *NDRG1* KO (magenta) and CNTR (blue) cells. Error bars are SEM corresponding to technical replicates. **k-l,** SW1990 and HPAC cell proliferation was analysed in 10% FBS using live image proliferation analysis in the presence of increasing concentrations of SGK1 inhibitor BLU6340 or vehicle control (DMSO). Error bars are SEM. No significant changes were observed. **m,** Fitted dose response curves for gemcitabine are shown in the presence of non-conditioned media (NC), CAF-CM, or CAF-CM supplemented with BLU6340 (250nM). Error bars are SEM corresponding to technical replicates. **n**, SW1990 and AsPC CNTR and *NDRG1* KO cells were treated with 1 µM gemcitabine (GEM), or 0.5 µM SN-38. SGK1 inhibitor (250 nM BLU6340) treated CNTR cells were also included in the analysis. 24h later the cells were fixed and stained for yH2AX followed by image acquisition and quantification by CellProfiler, data are shown as foci counts per single nucleus. >220 nuclei were analysed per condition. Mann-Whitney test was applied, magenta bars represent median. **o-r,** Cell cycle analyses of SW1990 and AsPC cells. **o, q** SW1990 and AsPC *NDRG1* KO and CNTR cells were subjected to double thymidine block and released into 10% DMEM. Time points were collected with unsynchronised (non-sync) cells, cells were fixed and processed for flow cytometry using PI for total DNA content, EdU-click coupled to Pacific Blue 450 azide as a marker for cells in S phase, phosphorylated histone H3 Ser10 (H3-pS10) conjugated to Phycoerythrin (PE) as a marker for mitosis. Due to partial emission spectra overlap between PI and PE it is possible to present total DNA content and mitotic population on the same axis. Representative cell cycle scatter plots showing propidium iodide (PI), phospho-histone H3 (pH3) (x-axis) and EdU-positive cells (y-axis) staining intensities. Plots were generated in Flowjo after gating on single cells only. Cells stained for each of the fluorophores were used as gating controls. Time point 0h shows synchronization of the cells at G1/S border. **p,r**, SW1990 and AsPC cells were subjected to double thymidine block, pre-treated with BLU6340 250nM for 16h and released into 10% DMEM supplemented with BLU6340 250nM when indicated. Time points were collected with unsynchronised (non-sync) cells, cells were fixed and processed for flow cytometry using PI for total DNA content, EdU-click coupled to Pacific Blue 450 azide as a marker for cells in S phase, phosphorylated histone H3 Ser10 (H3-pS10) conjugated to Phycoerythrin (PE) as a marker for mitosis. Representative cell cycle scatter plots showing propidium iodide (PI), phospho-histone H3 (pH3) (x-axis) and EdU-positive cells (y-axis) staining intensities. Plots were generated in Flowjo after gating on single cells only. Cells stained for each of the fluorophores were used as gating controls. Time point 0h shows synchronization of the cells at G1/S border. **s,** 2×10^6^ SW1990 CNTR and a mix of *NDRG1* KO clonal cell lines (KO mix) were injected into flanks of male athymic nude mice (10 animals/group). Representative images of tumours of CNTR versus *NDRG1* KO cells are shown; NDRG1 expression was evaluated by IHC, scale bar: 1mm (whole size tumour images), and 100 μm for magnified areas. **t,** Growth dynamics of SW1990 tumour xenografts based on tumour volume, error bars are SEM, some error bars are not visible due to small variation in sample group. Student’s t-test was applied. **u,** Tumour weights between CNTR and *NDRG1* KO groups at end point were plotted, student’s t-test was applied for significance, error bars are SEM. **v,** *NDRG1* was knocked down in AsPC cells using CRISPR/Cas9 and after selection, batch KD population and CNTR cells were pooled and injected to nude mice. Tumours were allowed to grow until palpable and treated with 25mg/kg gemcitabine 2× week for 13 days. Tumours were harvested and imaged. **w,** AsPC tumour volumes were measured at end point and plotted. Student’s t-test was applied for significance, error bars are SEM. **x,** AsPC *NDRG1* KD and CNTR IHC sections of the tumours were stained for NDRG1 expression and imaged. Scale bar: 2 mm. **y**, AsPC tumour weights were measured and plotted. Student’s t-test was applied for significance, error bars are SEM. **z,** AsPC xenograft tumours sections were stained for pan-cytokeratin to define cancer cells within the tumour, for Ki67 to define proliferative cells, and the nuclei were counterstained by DAPI. Whole tumour image analysis was performed in QuPath by applying appropriate thresholds and thus defining Ki-67 positivity. Percent of the Ki67 positive cells in relation to pan-cytokeratin positive cells within each tumour is plotted. Student’s t-test was applied for significance. Error bars are SEM.

**Extended Data Fig. 5.**
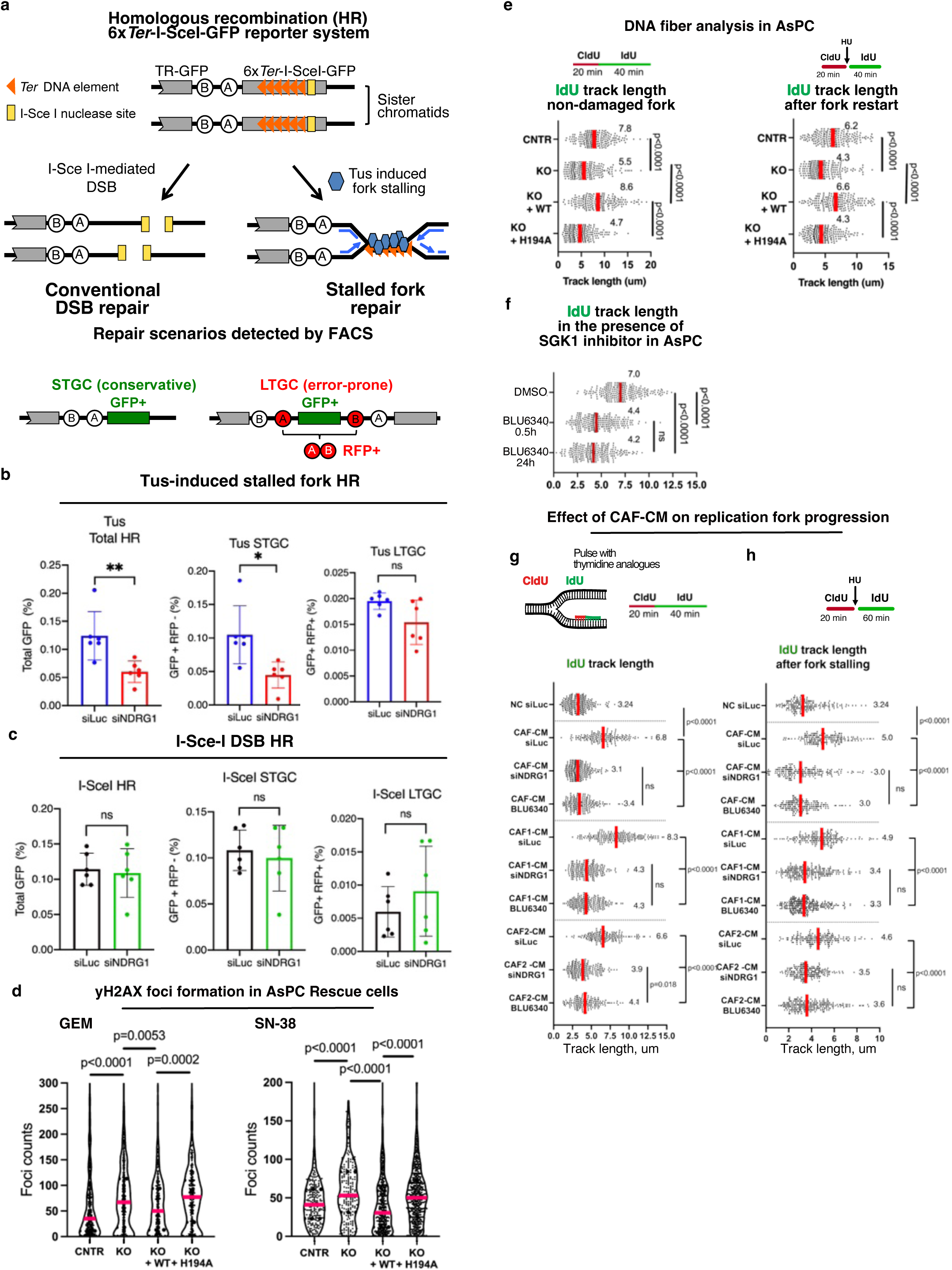
Loss of NDRG1 leads to replication stress and impaired stalled fork recovery. **a**, Schematic of the Tus/*Ter* replication fork barrier system allowing induction of site-specific replication fork stalling and measurement of homologous recombination (HR) at a defined locus in mouse embryonic stem cells. HR may be measured following either expression of the I-SceI nuclease and induction of a conventional double-strand break (DSB), or after site-specific replication fork stalling upon expression of the bacterial protein Tus. Tus binding to the 6× *Ter* array imposes a physical barrier to replication fork progression, stalling approaching replication forks, and triggering HR. (Orange triangle: 6×Ter element array. Blue hexagon: Tus protein, yellow rectangle: I-Sce-I endonuclease cut site). HR outcomes are quantified by flow cytometry: the reporter contains two non-functional heteroalleles of GFP (grey boxes) flanking synthetic exons encoding RFP in a tail to head orientation (A and B: 5’ and 3’ artificial RFP exons). HR may reconstitute a functional GFP allele and thus GFP+ cells - the 6×*Ter*-I-SceI GFP allele may undergo gene conversion upon copying a short region of the 5’-truncated GFP allele off the intact sister-chromatid, producing a wild-type GFP allele. These short-tract gene conversion (STGC) events represent conservative HR outcomes. A distinct minority of error-prone HR outcomes, termed long-tract gene conversion (LTGC), results in functional GFP with the duplication of the interior of the cassette harboring synthetic RFP exons. LTGC events produce cells that are both GFP+ and RFP+. **b-c**, Quantification of repair outcomes induced by fork stalling or I-*Sce-*I incision. Reporter cells were co-transfected with either empty vector, Tus or I-*Sce*-I endonuclease vectors and with siRNAs targeting either NDRG1 or control Luc. Transfected cells were analysed after 72 hours. Normalized STGC and LTGC repair frequencies were calculated and corrected for transfection efficiency. Triplicate samples and mean values from six independent experiments and Student’s t-test are shown. NDRG1 knockdown has a specific effect on HR at stalled forks (A); loss of NDRG1 inhibits total HR and error-free STGC at stalled forks (top panel), whereas it did not influence conventional DSB repair (I-*Sce*-I, lower panel). P-value: *<0.05, **,<0.01. ns: not significant. **d**, Quantification of yH2AX formation in AsPC CNTR, NDRG1 KO, and rescue (WT, H194A) cells after 1µM gemcitabine or 0.5µM SN-38 treatment (24h). yH2AX foci quantification was performed using CellProfiler, foci numbers per single nucleus were plotted and Mann-Whitney test was used for significance. Magenta lines represent median values. >200 cells were quantified per experiment. **e**, Replication fork dynamics in AsPC CNTR, *NDRG1* KO, and rescue cells (WT and H194A) were analysed by DNA Fiber Assay. Left panel: Cells were labelled with CldU followed by IdU as indicated. Right panel: Fork stalling was investigated by including HU (2mM, 1h) in between the CldU and the IdU pulse. Median IdU track lengths (µm) of >200 double-labelled fibers were analysed and shown in magenta. Mann–Whitney test was applied to test significance. **f**, To analyse the effect of short-term and long-term SGK1 inhibition on replication fork dynamics AsPC cells were treated with either vehicle control (DMSO), 250nM BLU6340 (0.5h) or pre-treated with BLU6340 for 24h, then labelled with CldU (20min) followed by IdU (40min). Median IdU track lengths (µm) of >200 double-labelled fibers are shown in magenta. Mann–Whitney test was applied to test significance. **g**, To assess the effect of CAF-CM on replication fork progression SW1990 cells (CNTR: siLUC, siNDRG1, and 250nM BLU6340 treated cells) were labelled with CldU (20min) followed by IdU (40min) in the presence of different CAF-CM media harvested from three patient lines supplemented with 2%FBS. Median IdU track lengths (µm) of >200 double-labelled DNA fibers are shown in magenta. **h,** To assess the effect of CAF-CM on fork stalling in SW1990 cells, experiment described in **(g)** was repeated with the inclusion of 2mM HU (1h) to stall the replication fork in between the CldU and IdU pulses. IdU pulse was 60 min. Median IdU track lengths (µm) of >200 double-labelled DNA fibers indicating fork restart are shown in magenta. Mann–Whitney test was applied to test significance.

**Extended Data Fig. 6.**
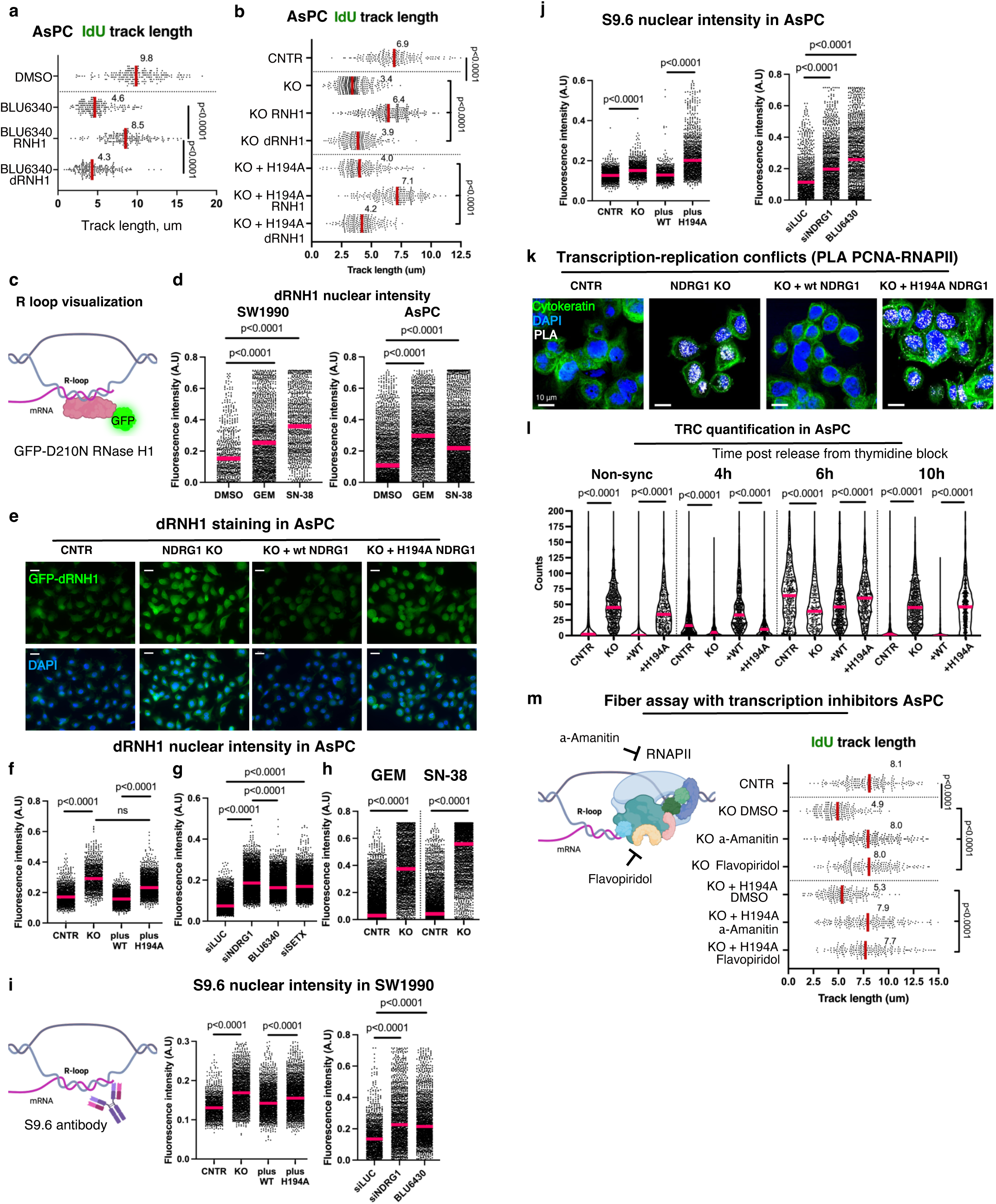
NDRG1 is involved in resolving R-loops. **a,** Replication fork progression in AsPC was analysed using DNA fiber assays. AsPC cells were treated with vehicle control (DMSO) or 250nM BLU6340 and co-transfected with WT-*RNase H1* (RNH1) or the catalytically dead *RNase H1* (dRNH1). Cells were pulsed with IdU (20min) followed by CldU (40min). Median IdU track lengths (µm) of >200 double-labelled fibers are shown in magenta. Mann–Whitney test was applied to test significance. **b,** Fork dynamics were analysed as in (**a**) using DNA fiber assays. AsPC CNTR, *NDRG1* KO and H194A cells were used and transfected with either WT-*RNH1*, or dead *RNH1* (dRNH1). Median IdU track lengths (µm) of >200 double-labelled fibers are shown in magenta. Mann–Whitney test was applied to test significance. **c,** Schematic of the recombinant RNase H1 construct used for R-loop visualisation. The bacterial purified GFP-labelled catalytic dead (D210N) RNase H1 can be used to probe RNA-DNA hybrids in fixed cells. **d,** SW1990 and AsPC cells were treated with vehicle control (DMSO), 1µM gemcitabine or 0.5µM SN-38 for 24h and analysed for RNA-DNA hybrid accumulation (R-loops) using the immunofluorescent GFP-dRNH1. Images were acquired, and dRNH1 nuclear intensity was analysed using CellProfiler, >670 cells were analysed per condition, Mann–Whitney test was applied to test significance, magenta lines are median values. **e,** dRNH1 immunofluorescence in CNTR, *NDRG1* KO and rescue (WT, H194A) AsPC cells. Scale bar: 20µm. **f,** Analysis of the fluorescence intensity of GFP-dRNase H1 in the nuclei of AsPC cells from **(e).** >950 cells were analysed per condition, Mann–Whitney test was applied to test significance, magenta lines are median values. **g,** GFP-dRNH1 nuclear intensity was analysed in AsPC CNTR cells, cells with NDRG1 knockdown (siNDRG1), or BLU6340 treatment. Knockdown of Senataxin (siSETX) was used a positive control to induce R-loops. At least 3000 nuclei were analysed per condition, magenta lines represent median, Mann-Whitney test was used for significance. **h,** To assess whether *NDRG1* KO increases R-loop content in the presence of chemotherapies AsPC CNTR and KO cells were treated with 0.5µM gemcitabine (GEM) or 0.5 µM SN-38 for 24h and analysed for RNA-DNA hybrid formation by GFP-dRNH1 staining. At least 1900 nuclei were analysed per condition, magenta lines represent median, Mann-Whitney test was used for significance. **i**,**j** S9.6 antibody that recognises RNA-DNA hybrids was used as an additional method to analyse R-loop formation in NDRG1-impaired cells. SW1990 and AsPC (either *NDRG1* KO-rescue panel or cells with NDRG1 knockdown/treated with BLU6340) were stained with S9.6 antibody. Nuclear fluorescence intensity of S9.6 staining was measured in CellProfiler, magenta lines represent median values, Mann-Whitney test was used for significance. >300 cells (SW1990) and >775 cells (AsPC) were analysed per condition. **k,** Representative images of the transcription-replication conflicts (TRC) in AsPC cells. Proximity ligation assay (PLA) between PCNA and RNA Pol-II (RNAPII) was performed in CNTR, *NDRG1* KO and rescue (WT, H194A) cells. PLA foci between PCNA and RNAPII are shown in white. Cytokeratin (green) was used to visualize cell shape, Scale bar: 10µm. **l,** SW1990 CNTR, *NDRG1* KO, and rescue (WT, H194A) cells were synchronized by double thymidine block at G1/S border and TRC formation was analysed at different time-points post dT release. TRC accumulation was also analysed in non-synchronized cells. TRCs were assessed by PLAs between PCNA and RNAPII, imaged and TRC foci counts analysed by CellProfiler. At least 200 nuclei per timepoint were analysed, magenta lines represent median, Mann-Whitney test was used for significance. At 6h all the lines had significant TRC load which were resolved in the CNTR and WT add-back cells at 10h timepoint. Note that KO and +H194A cells failed to resolve TRCs at the 10h timepoint. **m,** DNA fiber assay in AsPC CNTR, *NDRG1* KO and H194A add-back cells in the presence of transcription inhibitors α-amanitin (10mg/ml) and flavopiridol (10µM). Transcription inhibitors reduce TRC conflicts, thus preventing R-loops accumulation and rescuing replication fork dynamics in NDRG1-deficient or H194A NDRG1 expressing cells. Cells were pulsed with CldU (20min) and IdU (40min). Median IdU track lengths (µm) of >200 double-labelled fibers are shown in magenta. Mann– Whitney test was applied to test significance.

